# A general theory of coherence between brain areas

**DOI:** 10.1101/2020.06.17.156190

**Authors:** Marius Schneider, Benjamin Dann, Swathi Sheshadri, Hansjörg Scherberger, Martin Vinck

## Abstract

What does neuronal coherence tell us about neuronal communication? Does coherence between field potentials (e.g. LFP, EEG, MEG) reflect spiking entrainment or coupling between oscillators? Is it a mechanism for communication between brain areas, or a byproduct of interareal connectivity and spectral power? We hypothesized that interareal coherence is explained by the fact that outputs from one cortical area give rise to synaptic inputs in the same brain area, and correlated synaptic inputs in another area. Our mathematical analysis demonstrates that coherence between a sending and receiving area is precisely predicted from only two parameters: Interareal connectivity and oscillatory synchronization in the sending area. This model predicts narrow-band coherence even in case of a flat transfer function and in the absence of spiking entrainment in a downstream area, and reproduces frequency-specific Granger-causality patterns between brain areas (gamma feedforward, beta feedback). In general, we find that Granger-causality between field potentials is dominated by oscillatory synchronization in the sending area, whereas spiking entrainment is determined by the resonant properties of the receiver. Our model accurately reproduces LFP-LFP beta-coherence between macaque areas 7B and F5 in the absence of beta phase-locking within area F5. Together, these findings uncover a precise mechanistic model of interareal coherence as a (by)product of connectivity and power.

## Introduction

The brain is a dynamical system that generates intelligent behavior through interactions between brain areas (Buzsáki, 2006; Miller and Wilson, 2008; Varela et al., 2001; Bressler, 1995; Engel et al., 2001; Singer and Gray, 1995; Nicolelis et al., 1995; Siegle et al., 2019; Fries, 2015). These interactions can be studied by measuring temporal correlations between electrophysiological signals from multiple brain areas (e.g. coherence, Granger-causality, cross-correlations). Electrophysiological signals are commonly distinguished into spike recordings and measures of population synaptic activity, e.g. LFP, ECoG, EEG, MEG (Nunez and Srinivasan, 2006; Pesaran et al., 2018; Buzsáki et al., 2012; Mitzdorf, 1985; Einevoll et al., 2013) (referred to as “field potentials”). Field potentials have obvious disadvantages as compared to spike recordings, such as: (i) Loss of spatial resolution, (ii) volume conduction, and (iii) the fact that synaptic potentials are a mixture of local and afferent inputs (Pesaran et al., 2018; Nunez and Srinivasan, 2006; Buzsáki and Schomburg, 2015; Einevoll et al., 2013; Buzsáki et al., 2012). Yet, they have clear advantages: (i) They can be recorded non-invasively or from the cortical surface, and (ii) they can uncover weak interareal interactions by pooling over synaptic potentials in a large volume.

Field potentials from different brain areas show coherent activity in various frequency bands (Buzsáki, 2006). Interareal coherence is correlated with several cognitive and behavioral factors (Grothe et al., 2012a; Gregoriou et al., 2009; Salazar et al., 2012a; Richter et al., 2018; Colgin et al., 2009; Buschman and Miller, 2007; Fries, 2015; Varela et al., 2001; Babapoor-Farrokhran et al., 2017; Phillips et al., 2014; Olcese et al., 2016; Montgomery and Buzsáki, 2007; Von Stein and Sarnthein, 2000; Bressler et al., 1993; Brunet et al., 2014). Furthermore, distinct frequency bands are thought to play specific roles in interareal communication. For example, gamma (30-80Hz) and alpha/beta frequencies (10-30Hz) have been related to feedforward and feedback corticocortical communication, respectively (Buschman and Miller, 2007; Bastos et al., 2015; van Kerkoerle et al., 2014; Richter et al., 2018; Bressler et al., 2006; Mejias et al., 2016). Yet, the unequivocal functional significance and causal interpretation of these findings remains to be established. Does interareal coherence itself have an influence on interareal communication? Or, is interareal coherence a byproduct of interareal connectivity, and hence communication (coherence through communication)?

The interpretation of interareal coherence between field potentials is fraught with many problems (Pesaran et al., 2018; Buzsáki and Schomburg, 2015; Nolte et al., 2004; Nunez and Srinivasan, 2006; Einevoll et al., 2013; Vinck et al., 2015, 2010). A well-known problem is the spatial spread of electromagnetic fields (volume conduction) (Sirota et al., 2008a; Nunez and Srinivasan, 2006; Pesaran et al., 2018; Vinck et al., 2016; Carmichael et al., 2019; Parabucki and Lampl, 2017). Here, we investigate another major problem, which we refer to as the *synaptic mixing problem*: In the normal LFP range (*<*80Hz), field potentials primarily reflect summed synaptic activity (transmembrane currents) in a neuronal population (Einevoll et al., 2013; Pesaran et al., 2018; Nunez and Srinivasan, 2006; Buzsáki et al., 2012). These synaptic potentials can be decomposed into two parts: (i) Synaptic inputs caused by spikes from neurons in the *same* brain area, and (ii) afferent synaptic inputs caused by spikes from neurons in *other* brain areas. Likewise, spiking activity in one brain area (A) can cause synaptic potentials in the *same* brain area (A), and highly correlated synaptic potentials in *another* brain area (B) at a delay. We refer to these effects as “synaptic mixing”. Consequently, electric signals measured in area A and area B may, in part, be delayed copies of the same underlying signal, which would trivially cause interareal coherence and Granger-causality (Pesaran et al., 2018; Buzsáki and Schomburg, 2015). Because synaptic transmission is not instantaneous, the synaptic mixing problem cannot be solved with techniques that address the volume conduction problem (Trongnetrpunya et al., 2016; Pesaran et al., 2018; Nolte et al., 2004; Haufe et al., 2012; Vinck et al., 2015, 2011). Here, we develop a general theory of the way in which synaptic mixing determines interareal coherence, using mathematical analysis, simulations of neuronal populations, and analysis of interareal recordings.

## Results

### Beta-coherence between areas F5 and 7B

We first analyzed neural data, in which two distant brain areas show clear beta-synchronization between LFP signals. Beta-synchronization is thought to be involved in motor preparation, maintenance of a cognitive state and top-down modulation (Richter et al., 2018; Buschman and Miller, 2007; Salazar et al., 2012a; Bastos et al., 2015; Engel et al., 2001; Scherberger et al., 2005). We recorded from subdivisions of the parietal (area 7B) and premotor (area F5) cortex, which are involved in tasks like the reaching and grasping of objects (Dann et al., 2016). Area F5 is one of the main projection-targets of area 7B. The 7B-to-F5 projection is strong and long-range, as area F5 lies several cm’s away from area 7B (Johnson et al., 1996; Luppino et al., 1999; Markov et al., 2014).

We recorded LFPs and spiking activity using two 32-channel flexible microelectrode arrays per area (Figure 1a). We observed a clear beta-peak (≈ 20 Hz) in the power spectrum of 7B LFPs (Figure 1b). To analyze how single units were synchronized with LFPs, we computed the unbiased spike-field PPC value, which is proportional to the squared spike-field coherence (Vinck et al., 2012). Consistent with the LFP beta-peak, 7B units showed significant spike-LFP phase-locking in the beta-frequency band (Figure 1c). Spike-LFP locking was close to zero for frequencies outside the beta-band (Figure 1c). We also found coherence between the two 7B electrode-grids (Figure 1f,h).

**Figure 1:**
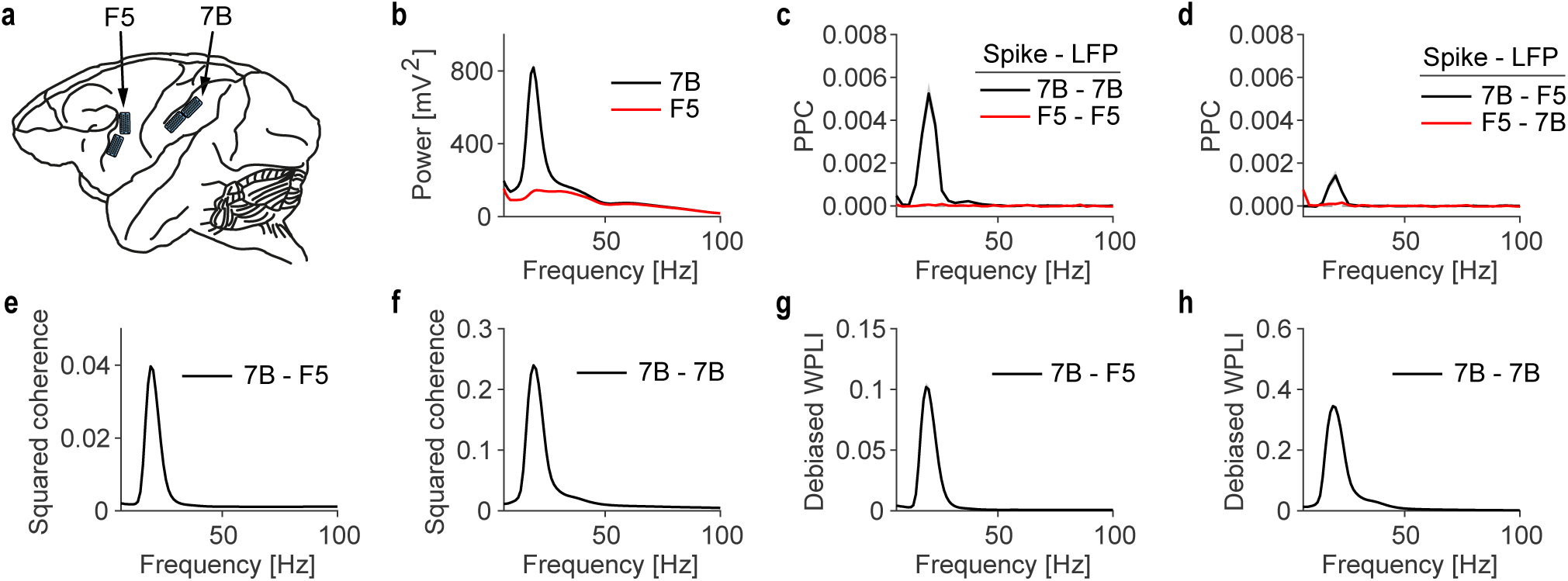
Coherence without spiking entrainment between macaque F5 and 7B **(a)** Illustration of electrode FMA grid recordings from the premotor areas F5 and 7B of a macaque monkey during grasping task (see Methods). We analyzed the memory period before motor execution. **(b)** LFP power spectrum for 7B (black) and F5 (red). **(c)** Spike-field phase locking (measured with PPC; see Methods) for different combinations of spikes and LFPs: Spikes in 7B to LFPs in 7B (black); spikes in F5 to LFPs in F5 (red). Phase locking values were averaged over all electrodes in corresponding grid. Spikes were pooled across all neurons in a session. The average number of neurons per session were 22.3 (7B) and 17.9 (F5). **(d)** Same as (c), but now between spikes in 7B to LFPs in F5 (black), and spikes in F5 to LFPs in 7B (red). **(e-f)** Coherence between medial and lateral 7B LFPs (f) and 7B and F5 LFPs (e). **(g-h)** Absolute value of weighted phase-lag index (Vinck et al., 2011) between 7B and F5 LFPs, as well as medial 7B and lateral 7B LFPs. The debiased WPLI is a measure of phase-synchronization robust to volume conduction (Vinck et al., 2011). All figures have standard errors of the mean. For PPC the standard error is across 4 conditions × 6 recording sessions. For coherence and WPLI, 6 sessions were pooled for the computation, and the standard error was computed across 4 conditions.

Next, we examined coherence between areas 7B and F5. We observed relatively strong and narrow-band beta-coherence between 7B and F5 LFPs (Figure 1e). This would suggest, *prima facie*, oscillatory coupling between these two areas. To rule out volume conduction, we computed the WPLI, a measure of synchronization that is not spuriously increased by volume conduction (see Methods) (Vinck et al., 2011). The WPLI spectrum showed beta-synchronization between 7B and F5 LFPs, suggesting that LFP-LFP coherence was not due to volume conduction (Figure 1g). This finding is consistent with the large spatial distance between 7B and F5.

Because of the clear beta-coherence between 7B and F5 LFPs, we expected to find beta oscillations in area F5, and interareal beta-synchronization between spikes and LFPs. Surprisingly, we did not detect significant beta-band spike-LFP phase-locking between F5 spikes and F5 or 7B LFPs (Figure 1c-d). The LFP power spectrum in area F5 was dominated by the 1*/f* component, and showed only a small peak in the beta-band (Figure 1b). Thus, there was clear beta-oscillatory activity in area 7B and beta-coherence between F5 and 7B LFPs, but no beta-synchronization within F5. How can this discrepancy be explained?

### Coherence predicted from connectivity and power

In this section, we will develop a model of interareal coherence between the field potentials in two areas (“Areas 1 and 2”). This model is applied to LFPs, but also applies to ECoG, EEG and MEG. We start with the case of unidirectional communication, where Area-1 projects to Area-2 with connection weight *w* (Figure 2a). In this case, Area-1 spikes can cause synaptic potentials both in Area-1 (through recurrent connections) and in Area-2. At the same time, Area-2 spikes will cause synaptic potentials in Area-2. Thus, the Area-2 LFP will be a mixture of synaptic inputs from Area-1 and Area-2. We model the measured signal in Area-1 as the sum of an oscillatory process and a broad-band process, e.g. 1*/f* pink noise. For now, we suppose that the intrinsic signal of Area-2 has no oscillatory component. The Area-2 LFP is therefore described as a linear mixture of its own background fluctuations and the synaptic inputs from Area-1.

**Figure 2:**
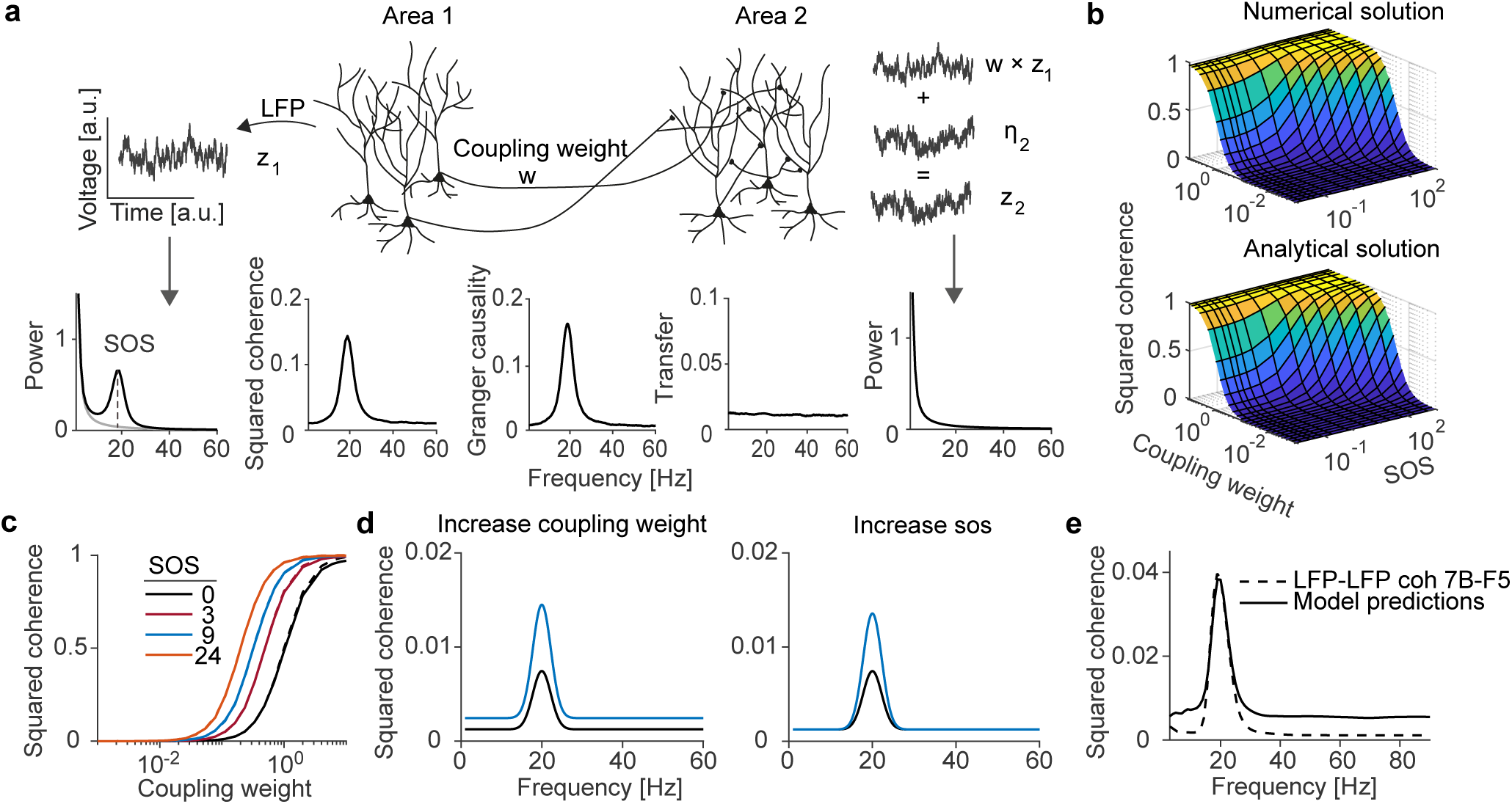
Interareal coherence as a (by)product of connectivity and power. **(a)** Illustration of the *synaptic mixing problem*. The Area-2 LFP is a linear superposition of intrinsic activity and afferent inputs from Area 1 weighted by *w*. The signal in Area-1 was modelled as a superposition of an oscillatory process and a 1*/f* background signal. The intrinsic activity in Area-2 was modelled by the same background signal. The power spectrum in Area-1 but not Area-2 shows a clear beta-peak. The coherence and Granger-causality spectra show clear beta peaks, but the transfer function is flat. The SOS (Sender Oscillation Strength) at the oscillatory frequency *f*_1_ = 20*Hz* was *SOS* (*f*_1_) = 14; *w* = 0.1. **(b-c)** Coherence as a function of the SOS and coupling weight. Analytical derivation matched the numerical simulations (c), performed using autoregressive models with varying oscillation strengths. Data: dashes. Model: solid. **(d)** Coherence spectra for two “behavioral” conditions, in which either the coupling weight (left) or the SOS (right) changed. The change in coherence was greater at the sender’s oscillation frequency. The parameters were: *SOS* = 10 (blue and black - left), *w* = 0.025 (black - left), *w* = 0.035 (blue - left) and *SOS* = 10 (black - left), *SOS* = 20 (blue - left), *w* = 0.025. Modelling interareal coherence between areas 7B and F5. Power spectra were fitted as a linear mixture of an AR(2) model with 1*/f* ^*n*^ background fluctuations (*w* = 0.069). The coherence at the peak frequency can be well reproduced, but that the coherence at other frequencies is not fitted well.

To understand the behavior of coherence in this model, we generated synthetic signals. Area-1 beta-oscillations were generated using dampened harmonic oscillators (AR(2); see Methods). The background processes were generated as 1*/f* pinknoise signals (see Methods). As expected, Area-1 showed a clear beta-frequency peak in the power spectrum (Figure 2a). However, because of the small connection weight (*w* = 0.069), there were no visible oscillations in the Area-2 power spectrum. Moreover, the transfer function from Area-1 to Area-2 was flat. This reflects the linear superposition of inputs from Area-1 onto Area-2, and the lack of any form of filtering. Despite this flat transfer function, coherence and Granger-causality showed clear spectral peaks in the beta-band. Thus, a narrow-band peak in the coherence spectrum emerged as a byproduct of synaptic mixing.

To generalize these observations, we derived an analytical expression for the coherence spectrum based on the connection weight *w* and the SOS (“Sender Oscillation Strength”). The SOS, *α*(*f*), is defined as the ratio of the power-spectral-density of the oscillatory component, *S* _11_(*f*), relative to the background signal, *H*_11_(*f*). The squared-magnitude coherence, which is approximately equal to Granger-causality, between Area-1 and Area-2 equals (see Methods)

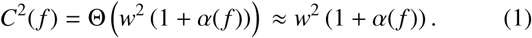

Here 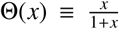 is a sigmoid-like function (Figure 2b). We confirmed this analytical expression by generating AR(2) and 1*/f* signals (Figure 2b,c). This analytical expression has two main implications:

1. The coherence peaks at the frequency where *α*(*f*) reaches a maximum. Even for small values of *w*, coherence can show a prominent peak if *α*(*f*) is large. In this model, band-limited coherence is a byproduct of communication: The coherence “itself” does not contribute to communication, because the transfer function is flat and there is no interaction between the inputs from Area-1 and the intrinsic activity of Area-2.
2. The dependence of coherence on coupling weight and SOS has a non-linear, sigmoidal form. Hence, changes in coupling weight cause steep changes in coherence for some values of *w*, but weakly affect the coherence for other values of *w*. Specifically, the derivative reaches a maximum for 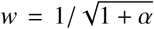. Thus, for large *α*(*f*), steep changes in coherence already occur for relatively small coupling weights. In general, the influence of the coupling weight depends on the value of the SOS. Furthermore, changes in the coherence can be caused both by changes in interareal connectivity and SOS (Figure 2d).

### Modelling the coherence between 7B and F5

We used the synaptic mixing model of coherence to reproduce the LFP-LFP coherence between areas 7B and F5. We modelled 7B beta oscillations as dampened harmonic oscillators, and the background processes as 1*/f* ^*n*^ spectra (see Methods). This model produced an LFP-LFP coherence spectrum with a clear beta-peak. However, it overestimated the LFP-LFP coherence at other frequencies (Figure 2e). Hence, our simple model of coherence cannot fully explain the interareal coherence between 7B and F5. Could there still be some “genuine” oscillatory coupling between areas 7B and F5?

### The influence of the projection-source coherence

In this Section, we uncover an additional mechanism that further suppresses the coherence related to non-oscillatory background fluctuations, namely the *projection-source coherence* (Figure 3a,b). In the model analyzed above, the Area-1 background fluctuations were projected onto Area-2 *in the same way* as the Area-1 oscillations. This assumes that the *inputs* into Area-2, caused by Area-1 activity, are a fully coherent copy of the total activity in Area 1.

**Figure 3:**
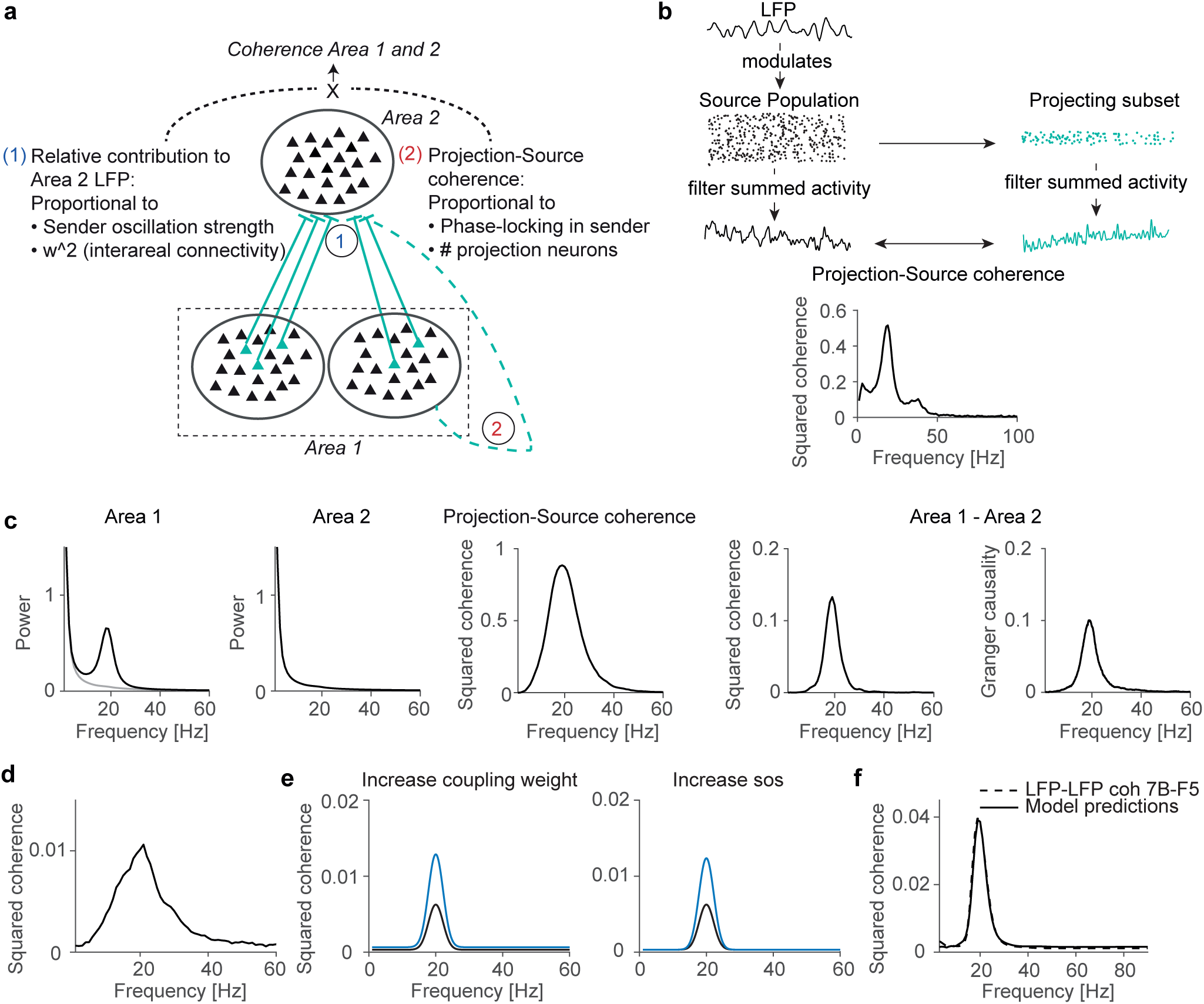
Interareal coherence is strongest at the frequency where the sender exhibits oscillatory synchronization. **(a)** Illustration of two mechanisms due to which coherence is suppressed at frequencies where the sender does not exhibit synchronization. The sending area consists out of multiple local sub-populations, whose projections may converge onto another area. We expect the high coherence at the oscillatory frequency, but low coherence for the background 1*/f* fluctuations. In addition, the sending area contains only a small population of Area-1-to-2 projecting neurons. The summed potential of these projecting neurons will be mostly coherent with the Area-1 LFP at the oscillation frequency. **(b)** Illustration of projection-source coherence. An LFP signal was generated as an oscillatory AR(2) process, and used to modulate the activity of 200000 neurons according to an inhomogeneous Poisson process. The average PPC peak between the neurons and the modulation signal was 0.007. A subset of 100 neurons represents the projecting neurons. The activity of the projecting and the source population were summed up and filtered with a 100HZ low pass filter. The resulting signals were used to calculate a projection source coherence. **(c)** Simulation where the subset of Area-1-to-2 projecting neurons is most coherent with the Area-1 LFP at the oscillation frequency. The interareal coherence and Granger-causality (right) show spectral peaks only at the oscillation frequency. In this case, the SOS at the oscillatory frequency was *SOS* = 14 and coupling weight *w* = 0.1; the oscillation was modelled as an AR(2). The background fluctuations in Area-1 were only partially transmitted, with a weight of 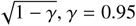 (see Methods). **(d)** Same as in Figure (c), but now with the presence of an oscillation in Area-2 with the same strength as Area-1. In this case the coherence is substantially lower than in (c), because the SOS equals 1 for all frequencies. However, it still exhibits a spectral peak because of the coherence of the Area-1 LFP with the Area-1-to-2 projection neurons. **(e)** Both increases in coupling weight and SOS cause a narrow-band increase in interareal coherence. The parameters were: *SOS* = 10 (blue and black - left), *w* = 0.025 (black - left), *w* = 0.036 (blue - left) and *SOS* = 10 (black - right), *SOS* = 20 (blue - right), *w* = 0.025. **(f)** LFP-LFP coherence between macaque 7B and F5 (dashed) can be reproduced (solid) by the synaptic mixing model shown in (c). The background fluctuations in Area-1 are only partially transmitted, with a weight of 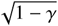, *γ* = 0.90, *w* = 0.077.

However, the Area-1-to-2 projections will originate from a relatively small subset of projection neurons (Markov et al., 2011, 2014; Han et al., 2018; Lur et al., 2016). The coherence between the summed activity of projection neurons and the Area-1 LFP, the *projection-source coherence*, may therefore be lower than 1. We will show that the projection-source coherence is an increasing function of two factors: (1) The fraction of projecting neurons; (2) the spike-LFP coherence of individual Area-1-to-Area-2 projecting neurons with the Area-1 LFP. In general, we expect that oscillations substantially enhance the spike-field coherence of individual neurons (Onorato et al., 2020; Sirota et al., 2008b; Vinck et al., 2016; Buffalo et al., 2011; Chalk et al., 2010; Buzsáki and Schomburg, 2015; Gregoriou et al., 2009)

A related point, is that the projecting area typically consists of spatially separated populations of projecting neurons. The outputs of these different populations may converge onto a single receiver. In the absence of coherent activity, the LFP tends to be highly local, spanning a small cortical volume (≈ 200*µ*m) (Lindén et al., 2011; Einevoll et al., 2013; Katzner et al., 2009). In this volume, the total number of neurons will be only around 160000*/*125 = 1280 neurons (for a density of 160000 neurons/mm^3^) (Christensen et al., 2007). Out of these, only few to tens of active neurons might be projecting to another given brain area. However, the coherence across populations can be strongly enhanced by rhythmic synchronization (Figure 1) (Buzsáki, 2006; Gray et al., 1989). This effectively increases the number of coherently firing projection neurons, and the spatial reach of the LFP (Lindén et al., 2011).

In the Methods section, we derive a general expression for the coherence between Area-1 and Area-2 as

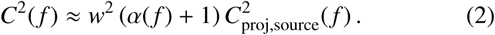

Here *α*(*f*) is the ratio of intrinsic power in the sender over intrinsic power in the receiver. This equation states that the original influence of the SOS and *w* on coherence is multiplied by the projection-source coherence. Thus, oscillations in the sender increase the interareal coherence because of two factors, *C*_proj,source_(*f*) and *α*(*f*).

Next, we derive an expression for *C*_proj,source_(*f*) (see Methods) as

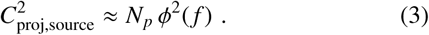

Here, *ϕ*^2^(*f*) is the squared spike-LFP coherence of an individual projecting neuron with the Area-1 LFP. The variable *N*_*p*_ represents the total number of Area-1-to-Area-2 projecting neurons. By combining all equations we state our main analytical result:

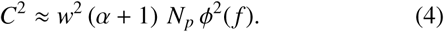

In sum, LFP-LFP coherence is determined by the following factors:

1. The connection weight *w relative* to other inputs into the receiver.
2. The number of active projecting neurons in Area-1, *N*_*p*_.
3. The spike-field coherence of projection neurons. From neural recordings, the value *ϕ*^2^(*f*) can be estimated using the unbiased spike-field PPC, which approximates the spike-field coherence (Vinck et al., 2012; Onorato et al., 2020; Vinck et al., 2016).
4. The sender-oscillation-strength *α*(*f*).

We further observe that the SOS and the spike-LFP PPC are proportional to each other, i.e. *α*(*f*) ∝*ϕ*^2^(*f*) (see Methods). For small values of *N*_*p*_, *α*(*f*) also depends linearly on *N*_*p*_ (see Methods). Moreover, the number of projecting neurons in an area should be proportional to the connection weight *w* between the areas (Markov et al., 2011, 2014). Hence we obtain the supra-linear relationship

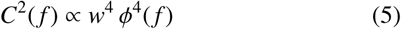

for small values of *w* and *ϕ*. Hence, 2-fold changes in spike-LFP coherence or firing rates in the sending area can, *ceteris paribus*, cause 16-fold changes in the squared interareal coherence. For larger values of *w* and *ϕ*, the expression takes the form *C*^2^(*f*) ∝ *w*^2^ *ϕ*^2^(*f*) as *C*_proj,source_ is bound by 1.

The connection weight depends on several factors: (i) The number of synaptic connections that are made into another area; (ii) factors modulating synaptic efficacy; (iii) the termination zone of the synapses, given that synaptic currents on basal and apical dendrites cancel each other out (Lindén et al., 2011); and (iv) the firing rates of the projecting neurons.

### Simulations of extended model

We validated our theoretical model with simulations. The projection-source coherence can be incorporated by simulating the activity in the sender as a weighted superposition of two 1*/f* background processes (see Methods): The projected 1*/f* background had a weight of 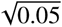, and the non-projected background a weight of 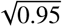. In this case, the projection-source coherence equalled 0.25 for the 1*/f* background process. As expected, the coherence was strongly suppressed at frequencies outside the oscillation frequency band (Figure 3c). Changes in interareal coherence due to changes in SOS and interareal conrnectivity also occurred in a narrow frequency-range, and were visible only at the oscillation frequency of the sender (Figure 3e).

An important consequence of the analytical expression in Eq. 2 is that a peak in the coherence spectrum can emerge even when the sender and receiving area have identical oscillation strength, in the absence of oscillatory coupling. By contrast, the simple synaptic-mixing model does not predict a peak in the coherence spectrum if *α*(*f*) equals 1 for all frequencies. However, due to its dependence on the projection-source coherence, the interareal coherence does attain a narrow-band structure (Figure 3d).

Finally, we confirmed the analytical expression obtained for the projection-source coherence (Eq. 3, 33, 39) with simulations (Figure 3b, 4a).

### Explaining coherence between areas 7B and F5

We observed that the coherence between 7B and F5 LFPs could not be reproduced by the basic synaptic mixing model (Figure 2e). We therefore applied the extended synaptic mixing model, in which the 1*/f* background is only partially transmitted. This model accurately reproduced the observed coherence between 7B and F5 LFPs (Figure 3f). We further modelled the interareal coherence based on the spike-LFP PPC within area 7B (Figure 4b). For this, we used the analytical expression for the coherence based on spike-LFP PPC, the SOS, the coupling weight and the number of projecting neurons. This model accurately reproduced the LFP-LFP coherence (Figure 4b). The required number of projecting neurons to reproduce the coherence was about 550. Note that for a given retrograde injection in F5, there are about two orders of magnitude more 7B-to-F5 projection neurons (Markov et al., 2011, 2014).

**Figure 4:**
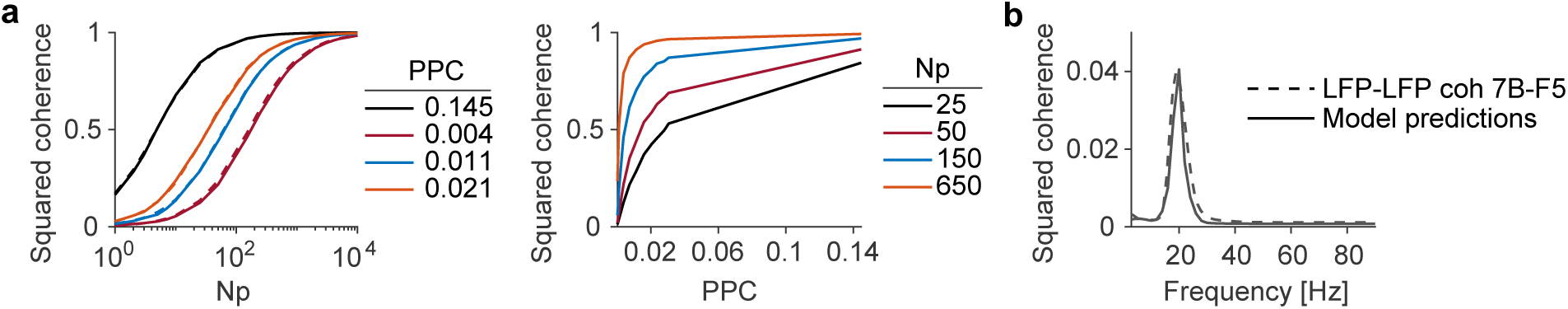
Analytical derivation of projection-source coherence, and interareal coherence predicted through spike-LFP Phase Locking in sender area **(a)** Squared coherence between Area-1 LFP and the summed activity of Area-1-to-2 projection neurons, as a function of the number of projecting neurons (Np) and the phase-locking of individual neurons (spike-field PPC). For this simulation, we generated an AR(2) signal in the sender and generated spikes in 10000 neurons according to inhomogenous Poisson processes modulated by this AR(2) process (see Methods; Figure 3b). The dashed lines show a tight match between our analytical derivations and the simulations (see Methods). **(b)** Interareal coherence spectrum between 7B and F5. Data: dashes. Model: solid. Model predictions were inferred from: the spike-LFP PPC within area 7B; the SOS *α*(*f*); and the coupling weight *w* of the model in Figure 2e. The total number of Neurons in area 7B is 100000 (which was arbitrarily chosen) and 550 of these are projecting to area F5.

### Explaining frequency-dependent interactions (gamma bottomup, beta top-down)

In the models presented above, we considered a unidirectional communication setting. However, brain areas are typically bidirectionally connected (Markov et al., 2011). The extent to which activity in Area-1 predicts activity in Area2, and vice versa, can be quantified using Granger-Gewekecausality (Geweke, 1982). Studies in primate visual and parietal areas suggest that *feedforward* and *feedback* Grangercausality are respectively strong at gamma and alpha/beta frequencies (Bressler et al., 2006; Bastos et al., 2015; van Kerkoerle et al., 2014; Buschman and Miller, 2007; Mejias et al., 2016; Michalareas et al., 2016; Richter et al., 2018). A possible interpretation of those findings is that brain areas communicate with different frequencies in the feedforward or feedback direction (Bastos et al., 2015). Alternatively, frequency-specific Granger-causality might result from the presence of distinct oscillation bands in different areas, not of frequency-specific transfer functions. In particular, oscillation frequencies and time constants decrease across the cortical hierarchy; gamma and beta oscillations are prominent in early visual areas and parietal cortex, respectively (see Figure 1b, (Bastos et al., 2015; Brovelli et al., 2004; Bosman et al., 2012; Vinck et al., 2016; Salazar et al., 2012a; Murray et al., 2014; Onorato et al., 2020; Spyropoulos et al., 2020; Scherberger et al., 2005).

We modelled the intrinsic signals in both Area-1 and Area-2 as the sum of a broad-band process (pink noise), and an oscillatory signal (Area-1: gamma; Area-2: beta) (Figure 5a). The areas were bidirectionally coupled. The synaptic mixing models accurately reproduced the Granger-spectra previously reported, with stronger feedforward and feedback Granger-causality at gamma and beta frequencies, respectively (Figure 5b-c). Thus, beta top-down and gamma bottom-up may be a trivial consequence of power gradients across the cortical hierarchy.

**Figure 5:**
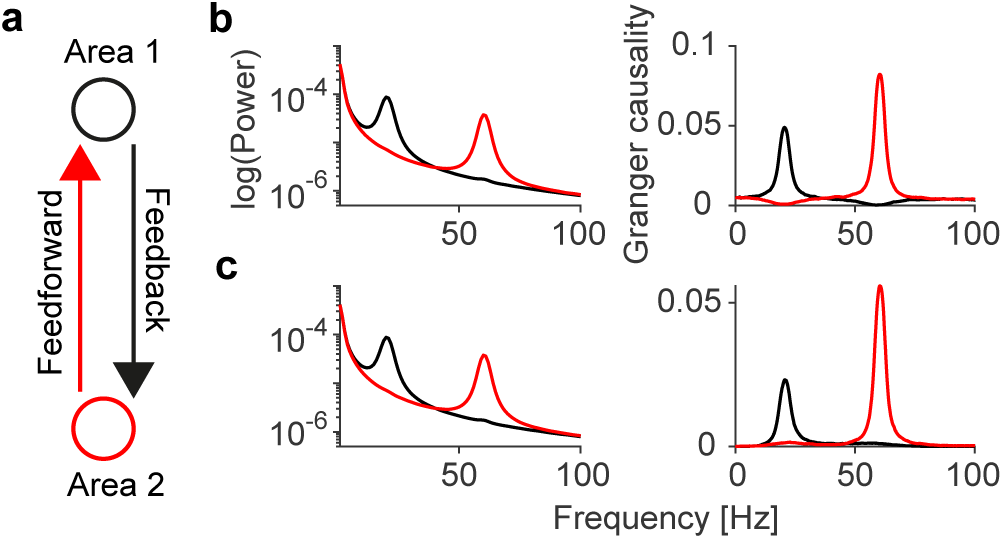
Gamma bottom-up and beta top-down Granger-causality are byproducts of connectivity and differences in power spectra. **(a)** We model feedforward (bottom-up) and feedback (top-down) interactions between two areas. The intrinsic activity of each area is modelled as a linear mixture of an AR(2) model and 1*/f* ^*n*^ background fluctuations. The AR(2) model of Area-1 is oscillating at 20 Hz. The AR(2) model of Area-2 is oscillating at 60 Hz. The signals are transmitted to the other area with a conduction delay of 5ms. **(b)** We simulate the interareal interactions according to the simple linear synaptic mixing model shown in Figure 2. Shown from left to right are: Power spectra of Area-1 and 2 and Granger-causality in feedforward (red) and feedback (black) direction. Thus, feedforward Granger at gamma and feedback Granger at beta frequencies can be reproduced by synaptic mixing alone, but some distortions are observed for feedback gamma and feedforward beta. **(c)** Same as (b), but now for the model where the background fluctuations in Area-1 are only parrtially transmitted, with a weight of 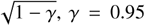. This again gives rise to gamma feedforward and beta feedback, but flattens the Granger-causality spectrum at frequencies outside the oscillation bands, producing a better fit to the published literature (Bastos et al., 2015). In (b) and (c) the signals were transmitted with a weight of *w* = 0.08.

### Synaptic mixing vs. spiking entrainment

In Figures 2-5, we showed how afferent synaptic inputs give rise to interareal LFP-LFP coherence due to synaptic mixing. However, afferent synaptic inputs from Area-1 can entrain the spiking output of Area-2 neurons (with spiking entrainment, we simply mean that afferent inputs change the probability of spike emission). The entrained spiking output of Area-2 neurons can in turn generate synaptic inputs in Area-2, which could further increase interareal LFP-LFP coherence (Figure 6a). Thus, LFPLFP coherence is determined by two factors, namely: (i) The coherence due to synaptic mixing, *C*_mixing_, and (ii) it is further increased by the coherence between the Area-1 LFP and the summed population spiking-activity in Area-2, *C*_entrainment_.

**Figure 6:**
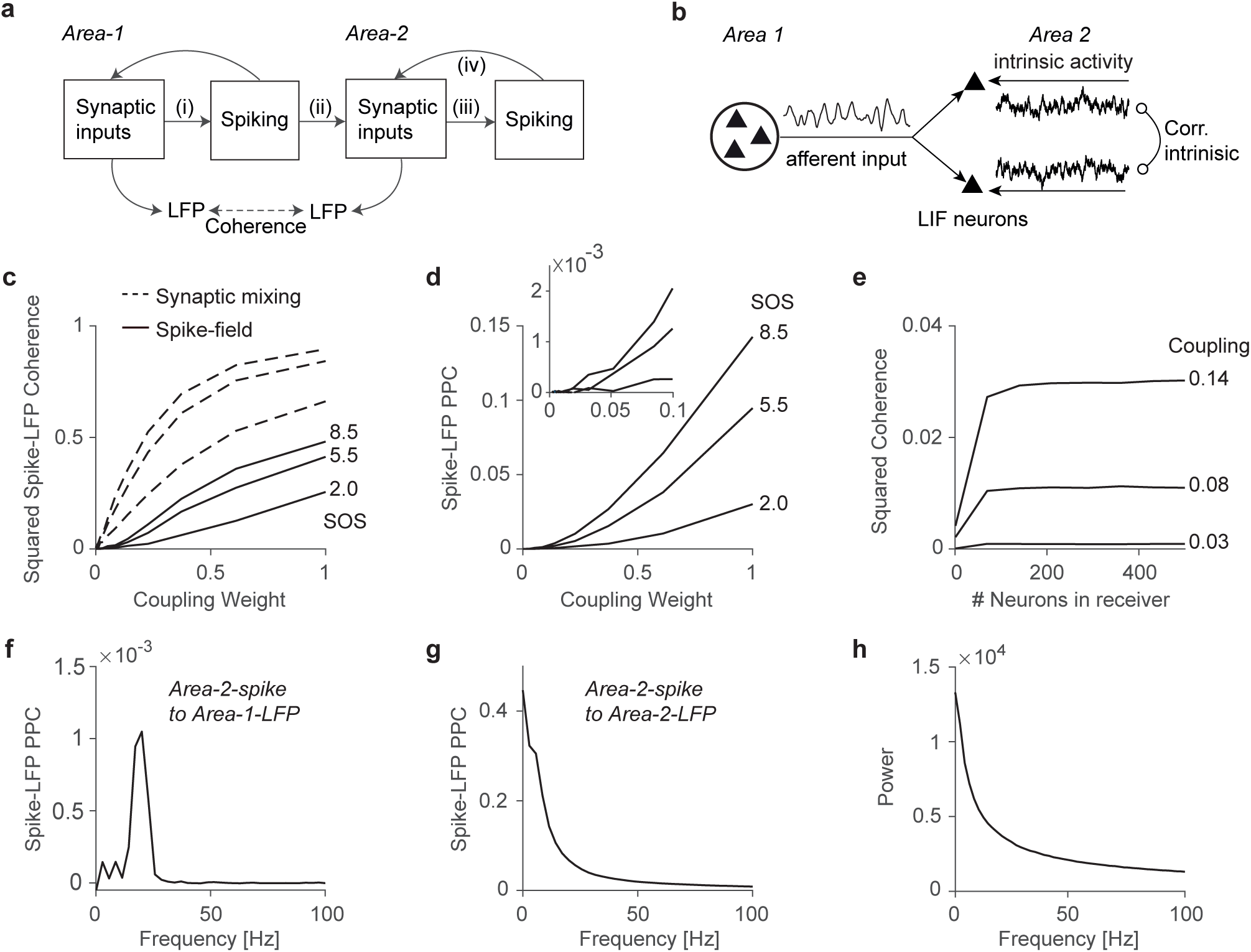
Dependence of coherence between Area-1 LFP and Area-2 spiking on connectivity, sender-oscillation-strength and number of neurons in Area 2. **(a)** Illustration of the different mechanisms causing coherence between two brain areas. Spiking activity in the sender (i) causes afferent synaptic inputs to Area-2 (ii). These inputs may not be a coherent copy of the LFP signal in Area-1 (projection-source coherence). Synaptic inputs in Area-2 give rise to additional spiking activity (iii), which through recurrent connections will give rise to synaptic inputs in Area-2 (iv). In case of resonance, oscillatory inputs may be amplified through recurrent dynamics at (iii) and (iv). The LFP, in the typical frequency range (*<* 80*Hz*), is a proxy of the population synaptic activity. **(b)** Illustration of model simulations on spiking entrainment in a receiver population. The receiver area is modeled as a population of leaky integrate-and-fire (LIF) neurons, whose intracellular potentials are a mixture of oscillatory input from sending Area-1 and intrinsic 1*/f* fluctuations in the receiver. The local fluctuations are correlated to different degrees. In the panels below, we use a correlation value of 0.5. **(c)** Squared coherence between summed population spiking activity in the receiver and the sender LFP, for different values of SOS and coupling weight (solid lines). LFP-LFP coherence due to synaptic mixing corresponds to dashed lines. Note similar dependences between LFP-spike and LFP-LFP coherence on SOS and coupling weight, but relatively higher LFP-LFP coherence due to synaptic mixing. **(d)** Spike-field phase-locking (of individual spikes; measured as PPC) between spikes in the receiver population and the LFP signal in the sender. Spike-field PPC shows similar dependencies on SOS and connectivity as spike-field coherence. Note relatively small spike-field PPC values for realistic connectivity weights around 0-0.1. **(e)** The number of neurons in the receiver determines the spike-LFP coherence. Inter-areal coherence between Area-1 LFP and Area-2 spikes increases with the number of receiver neurons. The SOS at the oscillatory frequency *f*_1_ = 20*Hz* was *SOS* (*f*_1_) = 5.5 **(f)** Spike-field PPC of receiver spikes to sender LFP. There is a clear beta-peak in the interareal spike-LFP PPC. **(g)** Spike-field PPC of receiver spikes to the receiver LFP. There is no beta-peak in the spike-LFP PPC in the receiver. **(h)** Power spectrum of the receiver LFP. Despite the afferent oscillatory input, the output of the LIF neurons shows no peak at the sender oscillation-frequency (beta). Instead, the spectrum follows the 1/f activity of the local fluctuations. (f to h) The parameters were: *SOS* = 5.5, *w* = 0.08.

Differences between *C*_entrainment_ and *C*_mixing_ will be primarily caused by the non-linear transformation of synaptic inputs into spiking outputs. An important question is whether the receiving circuit exhibits resonance in the frequency-band at which the sender oscillates. If it does, then the afferent oscillatory inputs can be amplified through recurrent interactions in the receiver, and *C*_entrainment_ may exceed *C*_mixing_ (we consider this case in Figure 7d). However, we first focus on non-linear transformations like sigmoids, in which case there is no network resonance.

**Figure 7:**
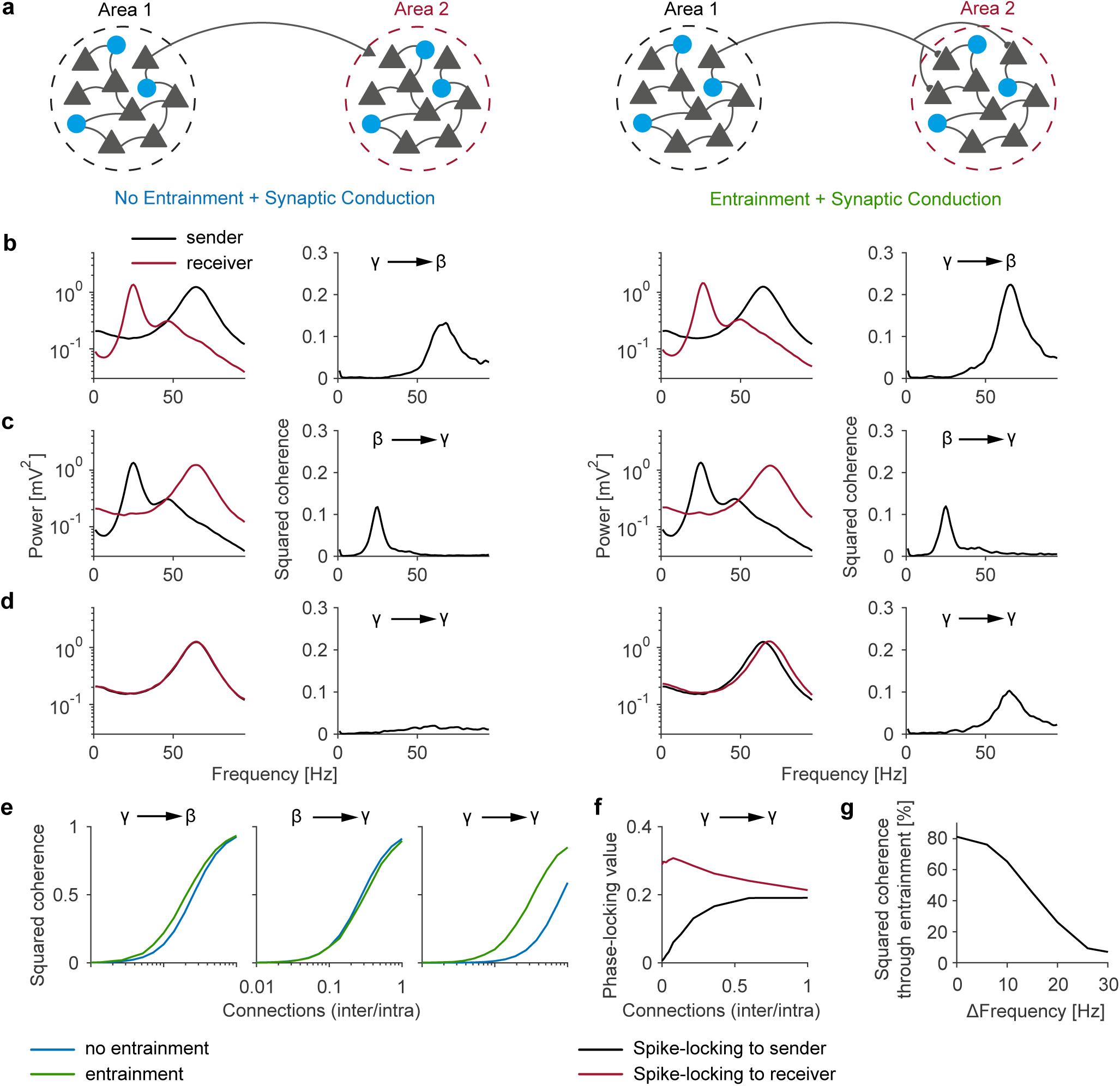
Spiking entrainment in the receiver does not substantially contribute to interareal LFP-LFP coherence, except when the frequencies in the sender and receiver overlap. **(a)** Illustration of the two models. Each area consisted of a population of spiking neurons whose dynamics were modelled by stochastic Wilson-Cowan equations. In the first model (left two columns), synaptic potentials due to inputs from Area-1 were superimposed onto the synaptic potentials from Area-2 itself. Neurons in Area-2 were “blind” to the synaptic inputs from Area-1, i.e. spiking entrainment was prohibited. The second model (right two columns) is identical to the first model, however synaptic inputs from Area-1 could now entrain the neurons in Area-2. **(b)** First two columns: Sender oscillates at gamma and the receiver at beta. Coherence spectra show clear peaks, following the power in the sender. Last two columns: Spiking entrainment increases coherence slightly. **(c)** and **(d)** Same as in (b), but now with different oscillation frequencies. When the oscillation frequency in the sender matches with the receiver, there is a increase in LFP-LFP coherence due to spiking (d). (b-d) all for coupling value of *w* = 0.1. **(e)** Change in coherence as a function of the ratio of inter- to intra-regional connection rates. **(f)** Spike-field phase-locking-value of neurons in the receiver population to the oscillations in the receiver (i.e. sum of all synaptic inputs caused by spikes in Area 2) and oscillations in the sender (i.e. sum of all synaptic inputs caused by spikes in Area 1). As the number of connections increases, the phase locking gradually increases. **(g)** Percentage of coherence explained by entrainment. Coherence spectra were calculated for 10 coupling values between 0.02 and The coherence peaks were integrated between *w* = 0.02 and *w* = 0.2.

We performed several simulations and analyses. First, we considered a scenario in which the receiver’s output is a sigmoidal transformation of its input and obtained analytical expressions for interareal coherence (see Methods). Second, we simulated a population of inhomogeneous Poisson neurons. These neurons were modulated by afferent beta-oscillatory inputs and intrinsic (i.e. local) 1*/f* fluctuations (see Methods; Figure S1). Third, we simulated a population of leaky integrate- and-fire (LIF) neurons (Figure 6b). In this case, individual LIF neurons received afferent beta-oscillatory inputs as well as intrinsic 1*/f* inputs that were correlated across neurons. Based on these simulations and analysis, we draw the following conclusions:

1. For all three simulations (sigmoid, Poisson, LIF), *C*_entrainment_ was lower than *C*_synaptic mixing_ (Figure 6c, S1c, S2a). One reason for this difference is that non-linear input-output transformations introduce a distortion in the coherence, and can introduce interactions between frequencies. Another reason is that the outputs of individual neurons are stochastic and sparse. Hence, spikes cannot encode afferent synaptic inputs without distortion. Distortion will be greater for a small number of neurons in the receiver and stronger correlations between local, intrinsic inputs in the receiver (Figure 6c,e, S1e, S2b).
2. If the population spiking-output in Area-2 is sigmoidally related to the summed synaptic inputs in Area-2, then *C*_entrainment_ will exhibit a similar dependence on the SOS, interareal connectivity and projection-source coherence as *C*_mixing_ (see Methods; S2c)). Also for Poisson and LIF neurons, we found that *C*_entrainment_ shows a similar, monotonic dependence on coupling weight and SOS as *C*_mixing_ (Figure 6c and S1). This also held true for the phase-locking of individual spikes to the LFP (spike-LFP PPC) (Figure 6d).
3. Narrrow-band phase-locking between receiver spikes and the sender LFP can occur in the absence of intrinsic oscillations in the receiver. In the LIF and Poisson simulations, the interareal spike-LFP PPC spectrum shows a clear beta-peak (Figure 6f). However, we did not find a concurrent beta-peak in the phase-locking between Area-2 spikes and the Area-2 LFP (Figure 6g). Accordingly, the receiver’s power spectrum did not show a beta-peak either (Figure 6h). Thus, rhythmicity in afferent inputs effectively disappeared in the receiver, both at the synaptic and spiking level.
4. In the absence of network resonance, we generally expect *C*_mixing_ to contribute more strongly to LFP-LFP coherence than *C*_entrainment_: First, note that the squared coherence measures the fractional part of power in the summed synaptic potentials in the receiver explained by the sender. Thus, the synaptic mixing component gives us the explained fraction of power at frequency *f* due to direct synaptic inputs from Area-1 (*C*_mixing_) (Figure 6a-ii). Because of the non-linear transformation from afferent synaptic inputs into spikes, the explained power in the summed population-spiking activity (*C*_entrainment_) will be lower than the explained power due to afferent synaptic inputs (*C*_mixing_) (Figure 6a-iii). Therefore, once the spikes in the receiver give rise to synaptic potentials through recurrent connections (Figure 6a-iv), they cannot explain a higher fraction of power in the synaptic activity than the afferent synaptic inputs themselves.

### E/I networks

To further investigate the contribution of spiking to LFP-LFP coherence, we modelled each area as a network of E/I neurons. Network dynamics were governed by stochastic Wilson-Cowan equations. These networks show noisy oscillations that mimic oscillatory behavior in the brain (Spyropoulos et al., 2020; Wallace et al., 2011; Powanwe and Longtin, 2019; Mejias et al., 2016). Note that the E/I networks did not contain dendritic lowpass filtering, which would have diminished the influence of E-E connections at higher frequencies (Buzsáki and Schomburg, 2015; Pike et al., 2000). We analyzed two types of scenarios:

#### 1. Mixing without entrainment

Area-1 spikes generated field potentials in Area-2, but Area-2 neurons were “blind” to the inputs from Area-1. Thus, there was synaptic mixing, but no spiking entrainment, and LFP-LFP coherence was strictly due to synaptic mixing.

#### 2. Mixing with entrainment

In addition to synaptic mixing, Area-2 neurons were modulated by synaptic inputs from the sender. We simulated different cases, e.g. gamma oscillations in the sender, beta oscillations in the receiver, or a combination of these. In Figure 7, we only show coherence; note however that Granger-causality is approximately equal to the squared coherence for unidirectional communication and yields qualitatively similar results (see Methods).

Without spiking entrainment, we found strong LFP-LFP coherence at the oscillation frequency of the sender (Figure 7b-c). However, in agreement with the results and arguments presented above, we found that spiking entrainment in the receiver did not substantially contribute to the interareal LFP-LFP coherence if the sender and the receiver had different oscillation frequencies (Figure 7b-c,e,g).

In Figure 7d, we considered a scenario where the sender and the receiver had the same oscillation frequency (gamma) and power. Without spiking entrainment, we found only weak gamma LFP-LFP coherence 7d. This is due to the fact that the SOS was now matched to the receiver oscillation-strength, i.e., *α*(*f*) = 1 for all *f* (see also Figure 3d). With spiking entrainment, we observed a strong increase in LFP-LFP gamma coherence. This reflects the resonant properties of the receiver, which amplify the afferent oscillatory input through recurrent dynamics. Overall, the relative contribution of spiking entrainment to LFP-LFP coherence decreased as a function of the difference in oscillation frequency between sender and receiver (Figure 7g). In the presence of resonant interactions, entrainment explained more than 50% of the squared coherence (Figure 7g). However, in the absence of resonance, entrainment only made a small contribution to the coherence (Figure 7g).

For all cases of sender-receiver frequencies, LFP-LFP coherence showed the stereotypical sigmoidal increase as a function of interareal connection strength. For the gamma-to-gamma simulation, phase-locking between Area-2 spikes and Area-1 LFPs was initially weak, indicating that most spikes in the receiver were triggered by its own intrinsic oscillations (Figure 7f). With an increase in interareal connectivity, we observed a gradual increase in spike-field phase-locking (Figure 7f). Thus, there was no sudden phase transition where the intrinsic oscillations in the receiver were fully phase-locked to the sender.

These simulations further demonstrate that LFP-LFP coherence and Granger-causality are of limited use for studying interareal interactions.

In particular, the frequency at which LFP-LFP coherence and Granger-causality show a peak is determined by the oscillatory properties of the sender, not the receiver (Figure 7). Furthermore, LFP-LFP coherence did not distinguish whether the sender-frequency matched the resonant-frequency of the receiver or not (compare *γ* to *β* with *γ* to *γ* in Figure 7e). However, the frequency at which spiking entrainment, i.e. actual information transfer, will be prominent, is determined by the resonant properties of the receiver, not the sender (Figure 8). Thus, if a lower brain area oscillates at gamma and the receiver resonates at beta, then LFP-LFP Granger-causality will show a clear gamma-peak (Bastos et al., 2015). However, one would expect that feedforward information transfer is specific to the beta-band, in which the receiver oscillates (Figure 8).

**Figure 8:**
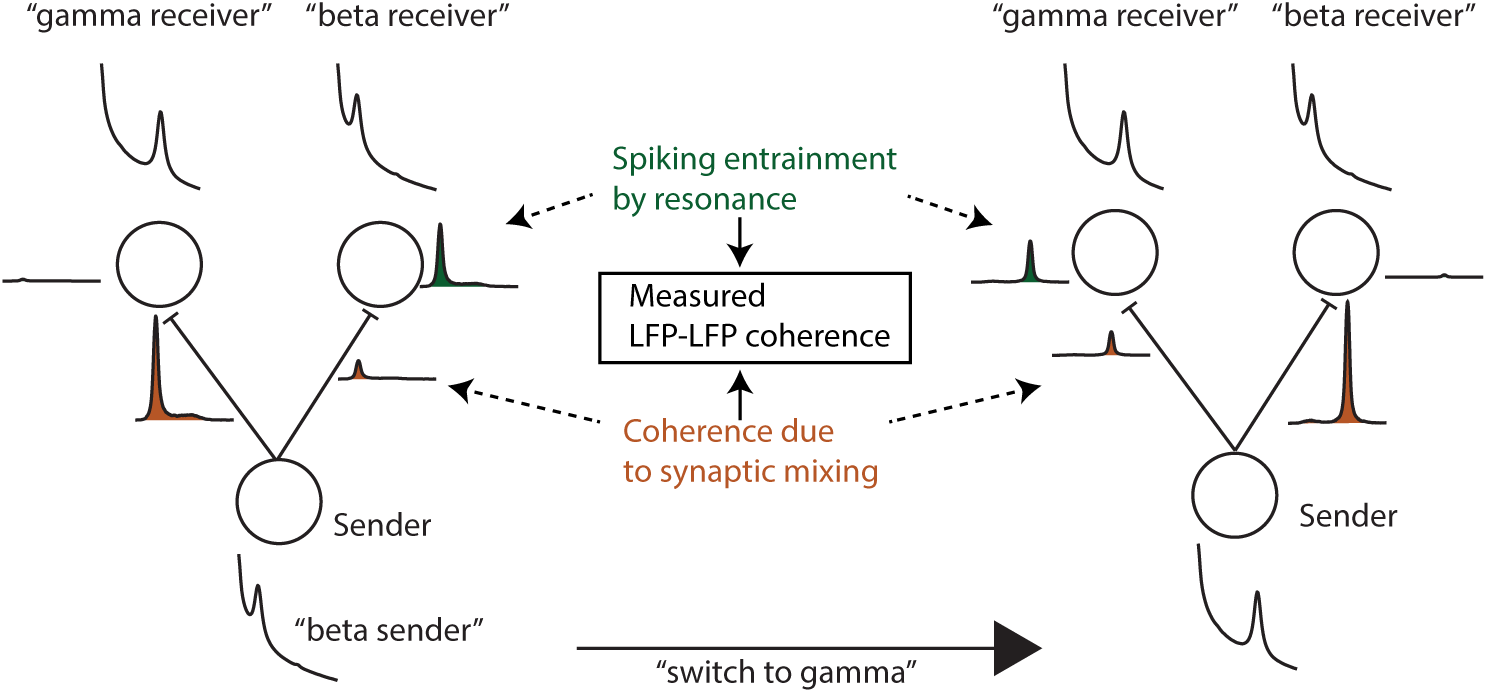
Illustration of difference between LFP-LFP coherence and spiking entrainment, and ability to switch communication by changing oscillations in the sender. In the left case, the sender oscillates at beta, and it would have a high LFP-LFP coherence with a receiver at gamma due to synaptic mixing. However, this LFP coherence does not translate into spiking entrainment. When the receiver also oscillates at beta, the LFP coherence due to synaptic mixing is lower, but due to resonance, the receiver will now exhibit more spiking entrainment. In the right case, the sender switches to gamma, and now switches communication to the gamma receiver. Paradoxically, LFP-LFP coherence might be higher between the sender and the receiver that communicate less.

## Discussion

The results of this paper demonstrate that interareal coherence between field potentials is a predicted byproduct of power and connectivity, and depends on four factors:

1. Spike-field coherence of projection neurons in the sender.
2. Oscillatory power in the sender relative to the intrinsic power in the receiver.
3. Interareal connectivity relative to other inputs into the receiver, which depends on the number of synapses, their strength, and the firing activity of neurons in the sender.
4. The number of active projection neurons.

Our results challenge conventional interpretations of interareal LFP coherence

### 1. Narrow-band interareal coherence does not indicate synchronization between intrinsic oscillations

Narrow-band coherence can simply arise from measuring a delayed copy of the sender’s oscillation in a receiving area, i.e. synaptic mixing. It does not require intrinsic oscillations in the receiver, in the form of a peak in the receiver’s power spectrum or local spike-field coherence.

### 2. Narrow-band coherence does not indicate that two areas predominantly communicate in the coherent frequency-band, and does not imply any specific transfer function

Due to synaptic mixing, narrow-band LFP coherence emerges even when the transfer function is flat and in the absence of any meaningful resonance or spiking entrainment in the receiver. In fact, even when information is transmitted at all frequencies, interareal LFP-LFP coherence may be visible only in a specific frequency band due to the specific nature of the LFP signal and the statistical properties of the coherence measure.

### 3. A narrow-band change in coherence does not demonstrate a change in communication or synchronization that is specific to the coherent frequency band

These narrow-band changes can simply result from changes in firing rates and phase-locking of projection neurons in the sending area. Furthermore, they can result from frequency-aspecific changes in effective interareal connectivity due to e.g. compartmentalized dendritic inhibition and neuromodulators (Batista-Brito et al., 2018; McGinley et al., 2015; Chiu et al., 2013).

Despite its limitations, interareal LFP-LFP coherence remains a useful tool for studying interareal connectivity and dynamic changes therein, especially in human ECoG. Our theoretical model of coherence and Granger-causality opens new avenues for mapping interareal connectivity in the human brain, providing an interesting alternative to diffusion tensor imaging, and leads to several practical suggestions:

1. Interareal connectivity can be most accurately estimated at the frequencies where the projecting area has a strong intrinsic oscillation (Figure 3 and 7).
2. LFP-LFP coherence carries little information about interareal connectivity in the absence of oscillatory synchronization (Figure 3). Thus, behavioral tasks should be used to stimulate a broad range of oscillations across cortical areas.
3. Using raw LFP coherence to predict connectivity leads to inaccurate results. Our analytical results show how LFP power differences should be factored in.
4. Given the dependence of interareal coherence on many factors, it is highly non-trivial to infer changes in interareal synaptic gain as a function of cognition or behavior. To make matters worse, controlling for the average firing rate or spike-field coherence is insufficient, because projection neurons are a highly specialized subclass of cells (Lur et al., 2016; Han et al., 2018).

### Spiking entrainment vs. synaptic mixing

We further analyzed the contribution of spiking to interareal LFP coherence. This yields two main conclusions:

1. In the absence of network resonance, synaptic mixing will make a stronger contribution to interareal LFP coherence than spiking entrainment (Figure 6, S1, S2, 7). In general, it is unclear to what extent afferent oscillatory inputs entrain spiking activity in the receiver (our Figure 1, Schomburg et al. (2014)). Entrainment depends on many factors, like non-linearities in single neurons and recurrent networks, low-pass filtering in pyramidal neurons (Buzsáki and Schomburg, 2015; Pike et al., 2000), and resonance (Figure 7, (Hahn et al., 2014)).
2. Narrow-band coherence between Area-2 spiking and Area-1 LFPs may simply reflect that the sender explains a large amount of variance in the frequency-band where the sender exhibits oscillatory synchronization. This spike-LFP coherence shows similar dependencies on SOS and connectivity as the coherence due to synaptic mixing. Importantly, narrow-band interareal spike-LFP coherence can occur in the absence of intrinsic oscillatory synchronization in the receiver (Figure 6): It is simply a consequence of connectivity/communication and does not demonstrate any functional consequence of coherence itself. Thus, the presence of narrow-band interareal spike-LFP coherence does not indicate that these frequencies have a privileged role in communication between brain areas. Furthermore, in the absence of resonance, oscillatory activity in the sender may not give rise to oscillatory activity in the receiver (Figure 6f-h). In this case, polysynaptic “synfire” chains are unlikely to occur, and the coherence across two brain areas will be proportional to the product of interareal connectivity values, 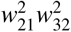. Consistent with this notion, experiments have shown that V1 activity correlates with spiking-activity in the V2 input layer (L4), but not with spiking activity in L2/3 of V2 (Zandvakili and Kohn, 2015).

### Communication-through-coherence or coherence-through-communication?

There is ample evidence that local synchronization is found in many cortical circuits and can increase firing rates in post-synaptic targets and boost plasticity formation (Kempter et al., 1998; Bernander et al., 1994; Salinas and Sejnowski, 2001; König et al., 1995; Abeles, 1982; Buzsáki and Draguhn, 2004; Vinck et al., 2013a; Sejnowski and Paulsen, 2006; Knoblich et al., 2010). A related but conceptually different idea is that *interareal phase-synchronization* is a mechanism for interareal communication (Varela et al., 2001; Bressler, 1995; Engel et al., 2001; Kreiter, 2006; Fries, 2005; Miller and Wilson, 2008; Bon-nefond et al., 2017; Palmigiano et al., 2017; Börgers and Kopell, 2008). Various aspects of these ideas have been summarized in the CTC (“communication-through-coherence”) hypothesis (Fries, 2005, 2009, 2015), which contains three premises:

1. Interareal coherence reflects phase synchronization between the *intrinsic* oscillations in the sender and receiver, e.g. due to entrainment or phase-resets by a common pacemaker.
2. Interareal communication is enhanced when afferent synaptic inputs consistently arrive at an excitable (“good”) phase of the intrinsic oscillation (Volgushev et al., 1998; Burchell et al., 1998).
3. Selective communication is implemented through selective coherence (Fries, 2015).

Our paper takes a different point of view: Two brain areas can only communicate if they are connected, and if they are connected, they will by default exhibit coherence at a “good” phase-relationship. This is due to the fact that the sending area will be coherent with the synaptic inputs that it causes in the receiving area, especially at the frequencies where the sender exhibits oscillatory synchronization. The resulting coherence is a consequence of communication, not a cause of it. To demonstrate coherence between *intrinsic* oscillations, it is therefore imperative to rule out that interareal coherence is not due to synaptic mixing. Otherwise, cause (connectivity and communication) can be easily confused with effect (coherence). If coherence is a byproduct of communication, then it is also unclear what experimental outcomes would possibly falsify CTC: If cognition is expected to increase interareal communication, e.g. due to attention, then an increase in coherence would be an *a priori* expected outcome.

CTC holds that communication between two connected areas is blocked by the absence of coherence (Fries, 2009). Yet, trivially, synaptic mixing models also predict that there is no coherence if there is no communication. Furthermore, communication is not consistently blocked by the absence of coherence, because interareal phase-relationships randomly fluctuate between “good” and “bad” phases (Akam and Kullmann, 2012). Alternatively, interareal communication may be prohibited by interareal coherence with a consistent “bad” phase-relationship (Volgushev et al., 1998; Burchell et al., 1998; Fries, 2005; Tiesinga and Sejnowski, 2010; Akam and Kullmann, 2012). This could block communication quite effectively (Volgushev et al., 1998; Burchell et al., 1998; Tiesinga and Sejnowski, 2010; Akam and Kullmann, 2012) and is not predicted by synaptic mixing models, but has not yet been demonstrated in empirical studies (Grothe et al., 2012a; Bosman et al., 2012). How can we disentangle coherence-through-communication from phase-synchronization between intrinsic oscillations? The strength of interareal coherence is a useful indicator: Because interareal connections are typically weak (Markov et al., 2014, 2011), synaptic mixing is unlikely to yield very high coherence values. These would suggest entrainment by a pacemaker, oscillatory coupling or volume conduction. However, correlational evidence for CTC consists of a moderate change in V1-V4 gamma-coherence with attention (from about 0.06 to 0.09, squared-magnitude coherence values below 0.01) (Ferro et al., 2020; Bosman et al., 2012; Grothe et al., 2012b). Notably, area V1 and V2 contain a very strong source of narrowband gamma (Roberts et al., 2013), which is associated with a unique class of excitatory neurons (Gray and McCormick, 1996; Onorato et al., 2020) and shows up to 300-fold power increases (Spyropoulos et al., 2020)). Thus, it is plausible that weak V1-V4 gamma coherence can be explained by synaptic mixing models (see Figure 5). A local increase in the firing rates and phase-locking of V1 or V2 projection neurons with attention would then be sufficient to increase V1-V4 coherence (Luck et al., 1997; van Kerkoerle et al., 2014; Buffalo et al., 2011; Chalk et al., 2010).

These considerations highlight a basic problem, namely how to experimentally identify *intrinsic* oscillations using LFPs, which contain a mixture of local and afferent synaptic inputs (Buzsáki and Schomburg, 2015; Pesaran et al., 2018; Saleem et al., 2017). The strength and prevalence of oscillations show great variation across the cortical sheet (Buzsáki, 2006). Several cortical regions exhibit very strong oscillations under specific sensory or behavioral conditions. For example: there is a strong source of gamma in V1/V2 (Gray et al., 1989; Peter et al., 2019; Vinck and Bosman, 2016; Onorato et al., 2020; Henrie and Shapley, 2005; Spyropoulos et al., 2020); beta in parieto-frontal cortex (Figure 1, (Scherberger et al., 2005; Dann et al., 2016; Brovelli et al., 2004; Salazar et al., 2012b; Hagan et al., 2012; Donoghue et al., 1998; Murthy and Fetz, 1996)); and theta and gamma in rodent hippocampus (Buzsáki, 2006; Colgin et al., 2009; Bragin et al., 1995). Coherence between these oscillatory sources and areas with weak or no intrinsic oscillations will be dominated by synaptic mixing (Figure 1, (Schomburg et al., 2014)). Due to synaptic mixing, oscillatory components will appear in areas without intrinsic oscillations. The contribution of afferent inputs to the LFP might depend strongly on the cortical layer: Feedforward projections target the granular layer 4, which has relatively little recurrent connectivity (Lund et al., 2003), and may not exhibit intrinsic oscillatory activity or resonance (Livingstone, 1996; Xing et al., 2012). Thus, layer-4 LFPs might be dominated by synaptic mixing of afferent inputs. Consistent with this notion, (Saleem et al., 2017) showed that in mouse V1, luminance/locomotion-related LFP oscillations in the 60-65Hz range are driven by LGN afferents and not affected by pharmacological suppression of V1 spiking activity.

Our main concern is the empirical evidence used to support theories of interareal phase-synchronization, not the theories themselves. Nevertheless, previous studies have raised numerous theoretical concerns that remain to be addressed, like the stochastic nature and strong stimulus-dependence of neocortical rhythms (Hermes et al., 2015; Ray and Maunsell, 2015; Burns et al., 2011), as well as the long time windows needed to distinguish competing inputs based on coherence (Akam and Kullmann, 2012). Our synaptic mixing model provides a straightforward explanation for narrow-band interareal coherence, indicating that it is a predicted byproduct of connectivity and oscillatory power, and depends on numerous factors that cannot be easily disentangled or controlled for. This prevents strong theoretical inferences based on empirical data and uncovers a circularity in theories like communication-through-coherence. It is unclear whether the synaptic mixing problem can be solved by using different connectivity metrics than Granger and coherence, for instance techniques from dynamical systems theory (Lowet et al., 2017). The problem likely needs to be addressed by detailed characterization of circuits at the micro-scale, using laminar current source densities and analysis of spiking (Buzsáki and Schomburg, 2015), and causal experiments (Saleem et al., 2017; Pesaran et al., 2018).

## Acknowledgements and Authorship contributions

We thank Prof. Dr. Wolf Singer, Dr. Georgios Spyropoulos, Patrick Jendritza, and Dr. Craig Richter for very helpful comments. Conceptualization: MS and MV. Mathematical analysis: MS and MV. Simulations and data analysis: MS. Macaque surgeries, recordings and data preprocessing: BD, SS and HS. Writing: MS and MV. Supervision: MV. This projected was supported by ERC Starting Grant to MV (SPATEMP) and a BMF Grant to MV (Bundesministerium fuer Bildung und Forschung, Computational Life Sciences, project BINDA, 031L0167).

## Methods

### Subjects

Neural activity was recorded simultaneously from many channels in one female rhesus macaque monkey (Animals S, body weight 9,7 kg). Detailed experimental procedures have been described previously (Dann et al., 2016). All procedures and animal care were in accordance with German and European law and were in agreement with the Guidelines for the Care and Use of Mammals in Neuroscience and Behavioral Research (National Research Council, 2003).

### Macaque data

The monkey was trained to perform a delayed grasping task. In this task, the monkey was either instructed to grasp a target with one of the two possible grip types (power and precision), or was free to choose between the grip types, as described in detail in previous studies (Dann et al., 2016). During instructed trials, the monkey was visually cued by one of two discs displayed on a monitor to perform the associated grip type. During free-choice trials, both discs were displayed, and the monkeys could freely choose between grip types. To encourage switching behavior during consecutive free-choice trials, the reward was iteratively reduced every time the monkey repeatedly chose the same grip type. Note that delayed-instructed trials were also part of the task. These trials were not analyzed in this study and are therefore not further explained. The monkey learned to perform the task with high accuracy of 95 +0.01 % SD successful trials on average.

Surgical procedures have been described in detail previously (Dann et al., 2016). In short, the monkey was implanted with four chronically implanted 32-channel microelectrode arrays (FMAs; Microprobes for Life Sciences; 32 electrodes; spacing between electrodes: 0.4mm; length: 1.5 to 7.1 mm monotonically increasing to target grey matter along the sulcus). Two arrays were positioned in part of the ventral premotor cortex (area F5) and two in area 7B, specifically around the anterior intraparietal area (AIP), yielding a total number of 128 channels. Electrode signals from the implanted arrays were amplified and digitally stored using a 128 channel recording system (Cerebus, Blackrock Microsystems; sampling rate 30 kS/s; 0.6-7500Hz band-pass hardware filter).

To detect spikes, electrode signals were first high-pass filtered with a median filter (window length 3ms) and then lowpass filtered with a non-causal Butterworth filter (5000Hz; 4th order). Next, common noise-sources were eliminated by applying principal component (PC) artifact cancellation, and spike waveforms were detected and semi-automatically sorted using ma modified version of the offline spike sorter Waveclus. Finally, redetection of the different average waveforms (templates) was used to detect overlaid waveforms. The exact procedures of spike detection are described previously (Dann et al., 2016). Only well-isolated single units were used for all analyses. To detect LFPs, electrode signals were first low-pass filtered with a median filter (window length 6.7 ms) and then high-pass filtered with a non-causal Butterworth filter (1 Hz; 4th order). In order to filter out line noise and their harmonics, additional band-stop filtering between 49 and 51 Hz and 98 and 102 was applied. Subsequently, signals were down-sampled from 30000 to 1000Hz by averaging 30 consecutive frames. Broken channels and trials containing movement noise were removed from all further data analyses. For this purpose, the total power, the correlation and the maximum deflection of all channels and trials were compared, and all outliers were discarded. Finally, to reduce the influence of the on-array ground and reference electrode on each array, the trimmed mean over all channels per array (leaving the highest two and the lowest two values per time point out) was removed by using linear regression. After spike and LFP detection, single-neuron spike events were binned in non-overlapping 1-ms windows to obtain an equal sampling rate of 1000 Hz for both signals. Subsequently, signals were aligned to cue and movement onset for the instructedand freechoice-task.

All analyses of macaque data were performed in Matlab (Mathworks) using custom scripts and the FieldTrip toolbox (Oostenveld et al., 2011). Power and coherence spectra were computed using integration windows of 0.35s, which were moved over the whole data in steps of 50ms. The epochs were Hann-tapered to avoid spectral leakage. The pairwise phase consistency (PPC) between spikes and LFPs was calculated using windows of 350ms around each spike (Vinck et al., 2012), using the spiketriggeredspectrum functions in the FieldTrip SPIKE toolbox. To compute the spike-LFP PPC, we first pooled the activity of single units in the area together, which gives the most sensitive estimate of entrainment in an area by increasing the number of pairwise phase comparisons (Vinck et al., 2013b).

### Predicting interareal coherence based on connectivity and power

In this Section, we derive an analytical expression for coherence based on interareal connectivity and power.

We start out from a unidirectional communication setting, where brain Area-1 projects to brain Area-2. The measured signals are denoted *z*_1_(*t*) and *z*_2_(*t*). In the following derivations, and our simulations, we assume that the signals are measured without the addition of extrinsic noise. That is we assume that all signals reflect neural activity, and we assume that there is no volume conduction.

We model the signal *z*_1_(*t*) in Area-1 as the sum of an oscillatory process *s*_1_(*t*) and a broad-band process, e.g. Pink noise, *η*_1_(*t*):

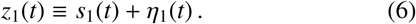

The intrinsic signal *z*_2_(*t*) of Area-2 has no rhythmic component and is modelled as a linear mixture of its own noise term and the projected input from Area-1,

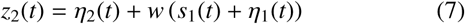

where *w* denotes the projection strength from Area-1 to Area-2. We assume that the background processes *η*_1_(*t*) and *η*_2_(*t* + *τ*) are linearly uncorrelated for all *τ*. For the purpose of mathematical derivation, we suppose that the power spectral densities of the broad-band processes are equal for all *f*, i.e. *H*_11_(*f*) = *H*_22_(*f*) ≡ *H*(*f*), with *f* frequency. We denote the spectral density of *s*_1_(*t*) as *S* _11_(*f*). We define the SOS (“Sender Oscillation Strength”) as

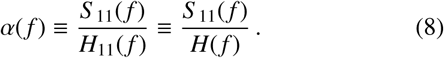

The cross-spectral density between *z*_1_(*t*) and *z*_2_(*t*) equals

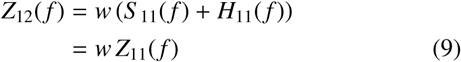

and is real-valued. Note that the other cross-terms fell out because we assumed that *η*_2_, *η*_1_ and *s*_1_ are uncorrelated. The squared coherence *C*^2^(*f*) between Area-1 and Area-2 is defined by

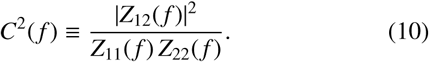

This simplifies as follows:

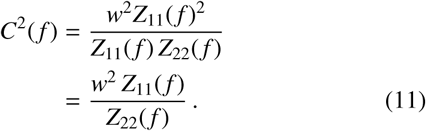

Since *η*_2_(*t*) and *s*_1_(*t*) are uncorrelated, we have

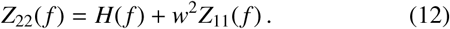

Eq. 8 now reduces to

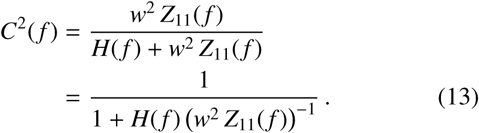

From Eq. 8 it follows that

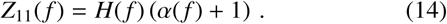

Hence *H*_22_(*f*) *w*^2^ *Z*_11_(*f*)^−1^ reduces to the expression

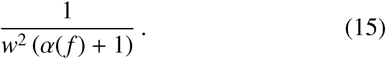

Thus the coherence can be simplified to

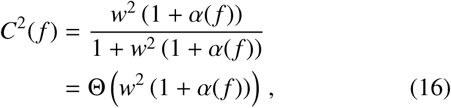

where 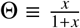 is the sigmoid function.

We can estimate the connectivity weight from the measurement variables by solving for *w* and *α*(*f*),

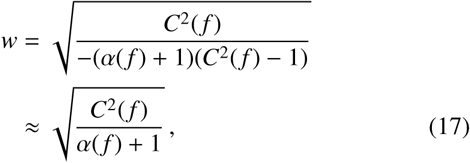

Here, the approximation is based on the first-order Taylor expansion of the coherence around *C*(*f*) = 0. We can also take the Taylor expansion around *w* = 0 for Eq. 16 and obtain

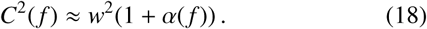

Note that the same model derivations (and the derivations below) pertain to Granger-causality, because for unidirectional coupling the following relationship holds between Geweke-Granger causality and coherence (Geweke, 1982):

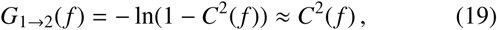

where the approximation was made based on the first order Taylor-expansion around *C*(*f*) = 0.

### Model of interareal coherence taking into account projection patterns

In the model above, we assumed that the signal received by Area-2 is fully coherent with the signal in the sender. As explained in more detail in the Results Section, this is likely not the case for two reasons: 1) The sender consists of subpopulations that are not fully coherent with each other, especially for frequencies where there is no oscillatory synchronization. 2) The number of projecting neurons in Area-1 may be small. Therefore, the coherence between the summed potential of Area-1-to-2 projection neurons and the Area-1 LFP (the projection-source coherence) may not be 1.

### Expression of the coherence based on power, interareal connectivity and the projection-source coherence

We first derive an expression of the interareal coherence that includes a linear dependence on the projection-source coherence.

We model the signals as

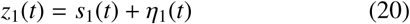

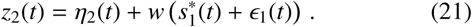

Here, 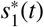 is the projected oscillatory signal into Area-2, and *ϵ* _1_(*t*) is the projected background signal into Area-2. The coherence between *η*_1_(*t*) and *E*_1_(*t*) is denoted *C*_*η,ϵ*_ (*f*). The coherence between *s*_1_(*t*) and 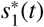 is denoted *C*_*s,s*_(*f*). We assume that *s*_1_(*t*) and 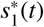 have the same power spectral densities. Likewise we assume that *η*_1_(*t*), *η*_2_(*t*) and *E*_1_(*t*) have the same power spectral densities.

We now obtain

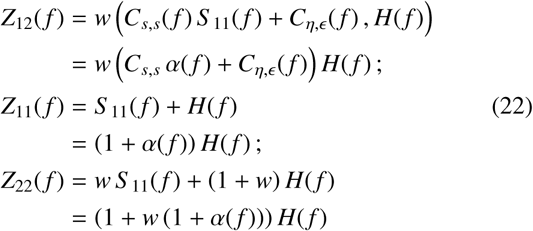

The squared coherence *C*^2^(*f*) now simplifies as

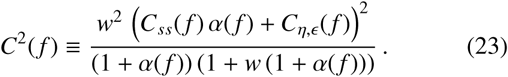

Plugging in *α*(*f*) = 0 for all *f* we obtain

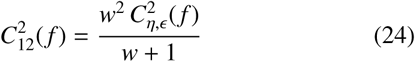

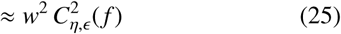

where the first-order Taylor expansion was made around *w* = 0. Thus, the squared coherence between areas scales with the coupling weight and the squared interareal coherence in the sender. For the oscillatory part, assuming the background fluctuations have coherence close to zero, we have

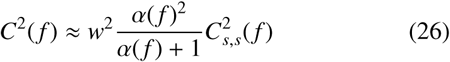

Following the same derivation, we can also obtain an expression for the squared coherence that combines both the noise and the oscillatory term as

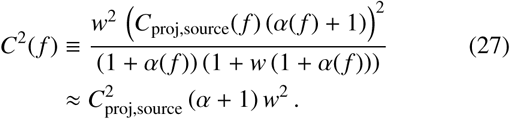

Here *C*_proj,source_ is the projection-source coherence. The variable *α*(*f*) is defined as the ratio of power of the intrinsic signal in the sender over the intrinsic signal in the receiver.

### Expression for the projection-source coherence

We derive the projection-source coherence based on *N*_*p*_ active (i.e. firing spikes) projecting neurons as follows. Let *x*_*i*_(*t*) be the activity of a single neuron in Area-1 with power spectral density *X*(*f*) for all *i*. The cross-spectral density of the *N*_*p*_ projecting neurons with the signal based on all *N*_*t*_ neurons in Area-1 equals

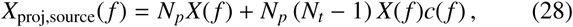

Here *c*(*f*) is the coherence between two individual neurons,

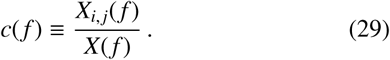

The factor (*N*_*t*_ − 1) accounts for the fact that each projecting neuron is fully coherent with itself. For simplicity, we assume that the cross-spectral density between any two neurons is real-valued (i.e. all neurons are on average coherent at zero-phase). The power of the signal in the source (Area-1) equals

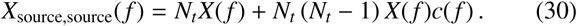

The power of the signal of the projection equals

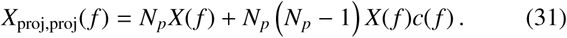

The squared coherence now equals

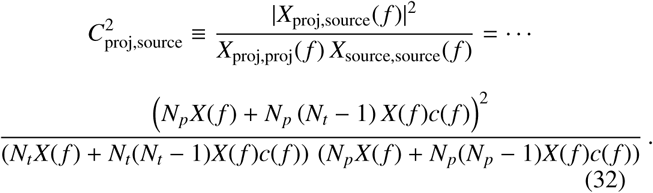

This simplifies further to

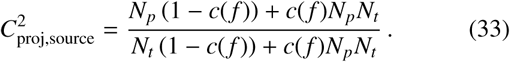

Plugging in 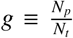, where *g* is the fraction of projecting neurons, we obtain

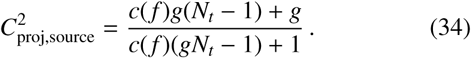

By taking the Taylor expansion around *c*(*f*) = 0, because the coherence between two individual neurons will be small, we obtain the first-order approximation

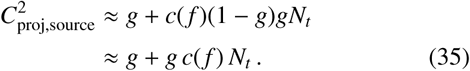

Here we removed the term (1 − *g*) because we can assume that *g* is typically close to zero. Hence the projection-source coherence is proportional to the fraction of projecting neurons, plus the coherence times the total number of projecting neurons. We can furthermore relate *c*(*f*) to the coherence of an individual neuron with the total signal in Area-1 (the spike field coherence). The squared-magnitude spike-field coherence can be expressed in terms of *c*(*f*) as

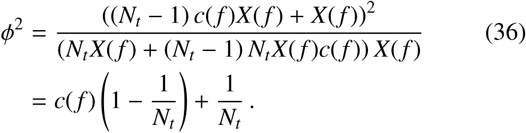

Note that we used

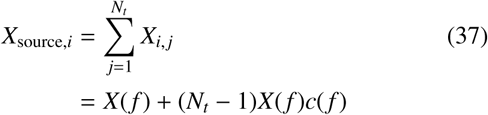

because we assumed all cross-spectra to be real-valued. Furthermore the total power in the source can be decomposed as

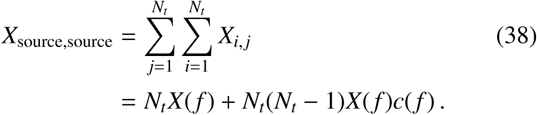

Solving Eq. 36 for *c*(*f*) yields

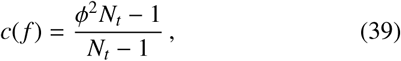

where *ϕ*^2^*N*_*t*_ ≥ 1. Plugging this into Eq. 35 we obtain the approximation

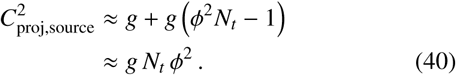

We thus obtain

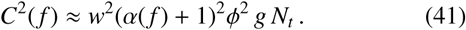

### Expression of the power on spike-field coherence

We further expect *α*(*f*) to be proportional to *ϕ*^2^: Let be *ϕ*(*f*) here is the consistency of single spikes (estimated by spike-field PPC) and divide the population into *N*_*t*_ spike trains of single spikes. The power due to the oscillation that is projected equals

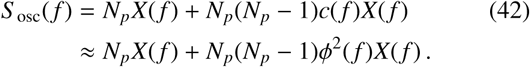

The power due to the background equals

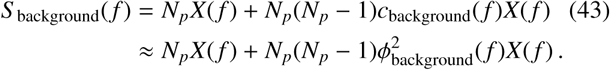

Here, *ϕ*_background_ is the spike-field coherence related to the background 1*/f* fluctuations, which may be non-zero. We note that if *N*_*p*_ is large enough, we have

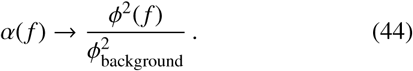

However, for small *N*_*p*_, we obtain the first-order Taylor expansion

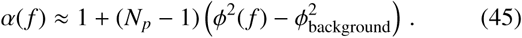

In this case, the SOS depends on *N*_*p*_. The reason for this dependence is that when *N*_*p*_ is small, the contribution of the phase consistency across neurons is relatively small and the intrinsic power due to the individual energy contributions weighs in.

### Non-linear dependence of coherence on spike-field coherence and connection weight

Because the connection weight *w* should be proportional to the total number of projection neurons (Markov et al., 2011), we therefore expect coherence to be proportional to *w* and *ϕ*. Combining all results we obtain:

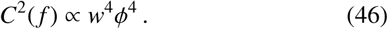

The factor *ϕ*^4^ follows from the dependence of *α* on *ϕ*^2^ and *C*_proj,source_(*f*) on *ϕ*^2^. The factor *w*^4^ follows from the dependence of *C*^2^(*f*) on *w*^2^, the dependence of *α*(*f*) on *N*_*p*_ and therefore *w*, and the dependence of *C*_proj,source_(*f*) on *N*_*p*_ and therefore *w*. When the number of projection neurons *N*_*p*_ and *ϕ*(*f*) is sufficiently high, the projection-source coherence *C*_proj,source_(*f*) should converge to one, and *α*(*f*) to 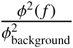. In that regime we obtain

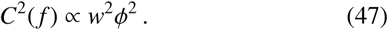

### Linear mixture with intrinsic noise in Area-1: simulations

For the purpose of simulations, we modelled the signal in Area-1 as follows:

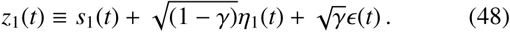

The intrinsic signal *z*_2_(*t*) of Area-2 is defined as a linear mixture of its own background fluctuations, and the input from Area-1:

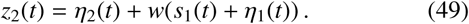

For the purpose of simulations, we assumed that the projected oscillatory component *s*_1_(*t*) is fully coherent with the oscillatory process in the sending area. We now obtain

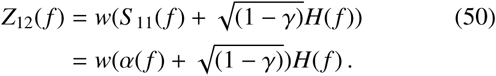

Since *η*_1_(*t*) and *γ*_1_(*t*) are uncorrelated:

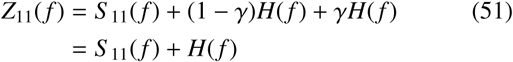

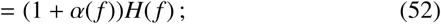

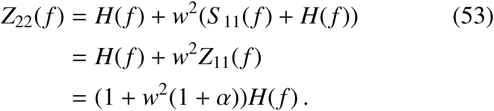

The squared coherence *C*_12_(*f*) now simplifies as

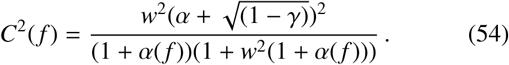

Plugging in *α*(*f*) = 0 we obtain

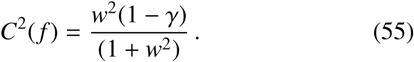

This is comparable to Eq. 24.

### Synthetic signals

#### Pink-noise signals

The background fluctuations in Figure 2 to 5 were simulated as 1*/f* ^2*/*3^ pink-noise processes. For every trial we generated a trace of white noise sample points. Each trace was Fourier transformed. The complex coefficients of the positive frequencies were multiplied by the 1*/f* ^2*/*3^-function. By concatenating the resulting coefficients with a flipped version of their complex conjugate, we obtained a spectrum following the 1*/f* ^2*/*3^-function. By inverse Fourier transforming the resulting spectrum we obtained a time series.

### AR(2) model

The oscillatory processes in Figure 2 to 5 were simulated using an AR(2)-model. The AR(2)-model is defined as

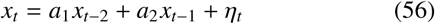

where *η*(*t*) is a white noise process with zero mean. To obtain a stationary signal, the roots must lie within the unit circle. If the AR process has complex conjugated roots it becomes a stochastic noise driven oscillator. The eigenvalues determine the strength of the oscillations.

### Wilson-Cowan model

In this section we summarize the population model from Figure 7. The Wilson Cowan Model is a stochastic network model of nonlinear neuron models. It is often used to demonstrate the appearance of oscillations on a network scale (Powanwe and Longtin, 2019; Wallace et al., 2011; Wilson and Cowan, 1972). Each area shown in Figure 7 was modeled by a Wilson-Cowan model, composed of fully connected *N*_*e*_ excitatory and *N*_*i*_ inhibitory neurons. The neurons were modeled as two-state Markov processes (one active and one quiescent state). In this model, the transition probability of the *i*th neuron to change from the active to the quiescent state is equal to

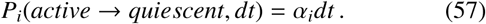

The transition probability of the *i*th neuron to change from the quiescent to the active state is equal to

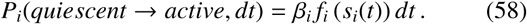

Here the activation function is defined:

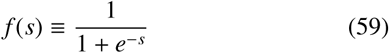

The total input current *s*_*E*_ to excitatory neurons and *s*_*I*_ to inhibitory neurons is defined:

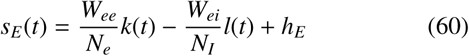

and

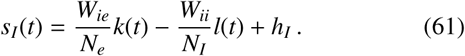

Here *h*_*I*_ and *h*_*E*_ are the external input current to the correspondent neuron types. The number of active excitatory neurons is referred to as *k*(*t*) and the number of active inhibitory neurons as *l*(*t*). The synaptic strength from excitatory neurons to inhibitory neurons is denoted *W*_*ie*_, and *W*_*ei*_ is the synaptic strength from inhibitory neurons to excitatory neurons. The total synaptic weight between excitatory neurons is referred to as *W*_*ee*_, whereas the total synaptic weight between inhibitory neurons is referred to as *W*_*ii*_.

The model determines the rates of transition between states by the variables *α* and *β*. However, since biological networks are stochastic processes, it is necessary to randomize the time of the next event. We achieved this by running the simulation with a Gillespie algorithm (Gillespie, 1977). In the scenario of “synaptic mixing with entrainment”, the excitatory neurons from Area-1 formed connections with the excitatory neurons of Area-2. This changed Eq. 60 for region 2 as follows:

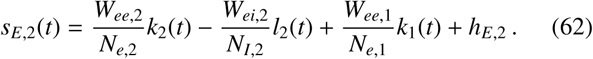

Whereas the neurons within an area were all-to-all connected, the inter-regional connection rate in 7b-d (right) was 10%. For simplification, each connection is represented in the LFP signal as one synapse. We calculated the LFP signal by convolving every incoming spike to an area with an alpha function 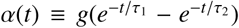. Here, the variable *t* is defined as the time relative to the spike onset and *α*(*t*) = 0 for *t <* 0. The factor *g* was equal to *g* = −1 for inhibitory synapses and *g* = 1 for excitatory synapses. Finally, the synaptic potentials of all input connections within an area were summed up to calculate an overall LFP signal of the corresponding area.

Each simulated area consisted of 800 excitatory and 200 inhibitory neurons. The neurons within one area were fully connected. In Figure 7 a-d, each neuron in Area-2 received inputs from 80 randomly chosen excitatory neurons within Area-1. All simulations in Figure 7 had the following parameter values, *W*_*ee*_ = 25.4, *W*_*ii*_ = 1.3, *W*_*ei*_ = 24.3, *W*_*ie*_ = 30, *h*_*E*_ = − 3.8, *h*_*I*_ = −3.8. Areas oscillating in the beta-frequency band had parameter values *α*_*e*_ = 0.038, *α*_*i*_ = 0.076, *β*_*e*_ = 0.379 *β*_*i*_ = 0.758. Areas oscillating in the gamma-frequency band had parameter values *α*_*e*_ = 0.1, *α*_*i*_ = 0.2, *β*_*e*_ = 1 *β*_*i*_ = 2.

### Extension to spiking activity

The same model developed for field-field coherence should apply to spiking activity, if spiking relates in a linear or sigmoidal way to synaptic inputs. Consider that *z*_2_(*t*) represents the average voltage fluctuations in the receiver. If population spiking activity is a linear function of *z*_2_(*t*), i.e. *y*_2_(*t*) = *z*_2_(*t*), then the same equation for coherence applies. Because spiking activity is stochastic and sparse for a single neuron, a population of neurons will contain additional variance that suppresses the coherence, i.e. we can write

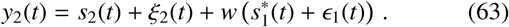

where *s*_2_(*t*) is the intrinsic signal in the receiver. This distortion *ξ*_2_, which should decrease with the number of neurons, will decrease the interareal spike-field coherence by increasing the intrinsic power in the receiver (see Figure 6 and S1). Next, consider the case where the population spiking activity is a standard sigmoidal activation function of *z*_2_(*t*), i.e.

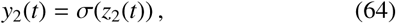

where *σ*(*x*) ≡1*/*(1+exp(−*x*)). In analogy to the data processing equality, we expected that the coherence after the transformation should always be lower than in the linear case, because the signal gets distorted by the sigmoid transformation, and coherence expresses the amount of variance that can be explained by linear prediction (see Figure S1 and S2). Assuming that *w* is relatively small, we use the Taylor-expansion around *w* = 0 and obtain

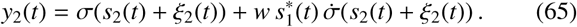

Here, 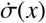 denotes the first derivative of the sigmoid function at *x*. Note that

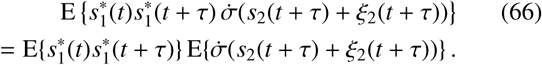

We can thus scale the signal as follows. Define a new transformation function by scaling inside the sigmoid as

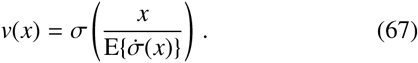

Assuming that *s*_1_(*t*) and *s*_2_(*t*) are statistically independent, we can see that the resulting coherence between *z*_1_(*t*) and *y*_2_(*t*) now equals

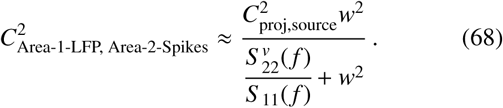

where 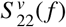 is the spectral density function of *v*(*s*_2_(*t*) + *ξ*_2_(*t*)). Here, we can recognize that the equation has the same form above and is scaled by the weight and the projection-source coherence. For small *w*, this dependence is linear. To conclude, in case of a sigmoid input-output curve:

1. For small *w*, the squared spike-field coherence scales with *w*^2^ and 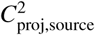, similar to the case of field-field coherence.
2. Coherence between Area-1 LFP and a population of spikes in Area-2 will be lower compared to field-field coherence because the spikes will be a noisy approximation of the input signal (for a finite population) (see Figures 6, S1, S2). Suppose that *y*_2_(*t*) reflects the superposition of spiking traces from a population of neurons. The number of neurons that we superimpose in the receiver has two effects: (i) If we assume that each neuron receives the same input, then adding more neurons increases the coherence between the Area-1 LFP and the Area-2 spikes, because the population sum becomes a more accurate approximation of the LFP (Zeitler et al., 2006; Vinck et al., 2012) (i.e. in the equation above, *ξ*(*t*) will decrease) (see Figures 6 and S1). (ii) Each neuron in the receiver may receive synaptic inputs from only a few Area-1 projection neurons. That is, the number of projection neurons that target a given neuron in the receiver will now be equal to *N*_*p*_*k*, where *k* is the fraction of projection neurons that projects to a single neuron in the receiver. Therefore, 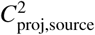 may be very small for a single neuron, and it should increase with number of neurons we superimpose. Thus, the coherence between the sum of a population of Area-2 neurons and the Area-1 LFP should depend in a non-linear way on the number of neurons we sum over.
3. We expect that if the spikes are a non-linear function of the input signal, there will be a frequency-dependent distortion in interareal coherence (see Figure 6, S1, S2). We expect this always to be a reduction if there is no network resonance, which remains to be proven. The distortion might be greater at higher frequencies, because the sigmoid transformation of low-frequency fluctuations can add noise to high frequencies.

### Poisson modulated neurons

In this section, we describe the simulations shown in Figure S1. To investigate the consequences of a sigmoidal input-output relationsip on interareal coherence, we simulated a population of in-homogeneous Poisson neurons. Area-2 was modelled as a population of up to 1000 neurons. The firing of the neurons was modulated by a synthetic LFP signal according to an inhomogeneous Poisson process. The modulation signal was a mixture of the afferent oscillatory inputs and the intrinsic 1*/f* fluctuations. This mixed modulation signal was passed through a sigmoid function and normalized on the standard deviation of the signal. We implemented the inhomogeneous poisson process by applying the timerescaling theorem (Brown et al., 2002). In the first step we generated spike times from a homogeneous Poisson process with unit firing rate. Thereupon we applied the inverse of the cumulative rate function to each event time (Nawrot et al., 2008). The average firing rate of the neurons across trials was 2Hz. There was no 1*/f* input from the sender area, i.e. the coherence followed the extended synaptic mixing model shown in Figure 3c-f.

### Leaky integrate and fire population

In this section, we describe the simulations shown in Figure 6. In order to investigate the effects of spiking entrainment on interareal coherence in a biologically realistic way, we modelled the receiver population as *N* leaky integrate-and-fire (LIF) neurons. Each neuron received an input current, which consisted of an afferent oscillatory signal *s*(*t*) and a local, intrinsic input *η*_*i*_(*t*). The local input *η*_*i*_ is modelled as a 1*/f* pink-noise process described before, and defined as

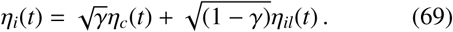

Here, *η*_*c*_(*t*) represents the shared fluctuations across receiver neurons, whereas *η*_*il*_(*t*) represents local fluctuations that are specific to neuron *i*. The parameter *γ* scales the temporal correlation between the neurons in the receiver population. The oscillatory signal *s*(*t*) was generated using the AR(2) model that we described above. We denote the power spectral density of the total background fluctuations in the receiver, 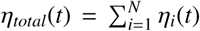, as *H*(*f*). The power spectral density of oscillatory input *s*(*t*) is defined as *S* (*f*). We define the SOS of a population of *N* neurons as

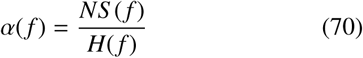

The dynamics of the membrane potential *V*_*m*_(*t*) of neuron *i* is defined by

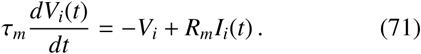

Here, *τ*_*m*_ is the membrane time constant, *R*_*m*_ the membrane resistance, and *I*(*t*) the input current.

The input current *I*_*i*_(*t*) of neuron *i* is defined as

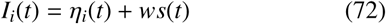

where *w* denotes the projection strength from Area-1 to Area-2. At *t* = 0 the membrane voltage is set to the resting potential *V*_*rest*_. Whenever the membrane potential passes a threshold *V*_*th*_, the neuron elicits a spike and the membrane potential is reset to *V*_*reset*_. The LIF neurons in Figure 6 and S2 had the following parameters: *τ*_*m*_ = 20*ms, V*_*rest*_ = − 60*mV, V*_*reset*_ = − 50*mV, V*_*th*_ = − 30*mV* and *R*_*m*_ = 100*M*Ω.

**Figure S1:**
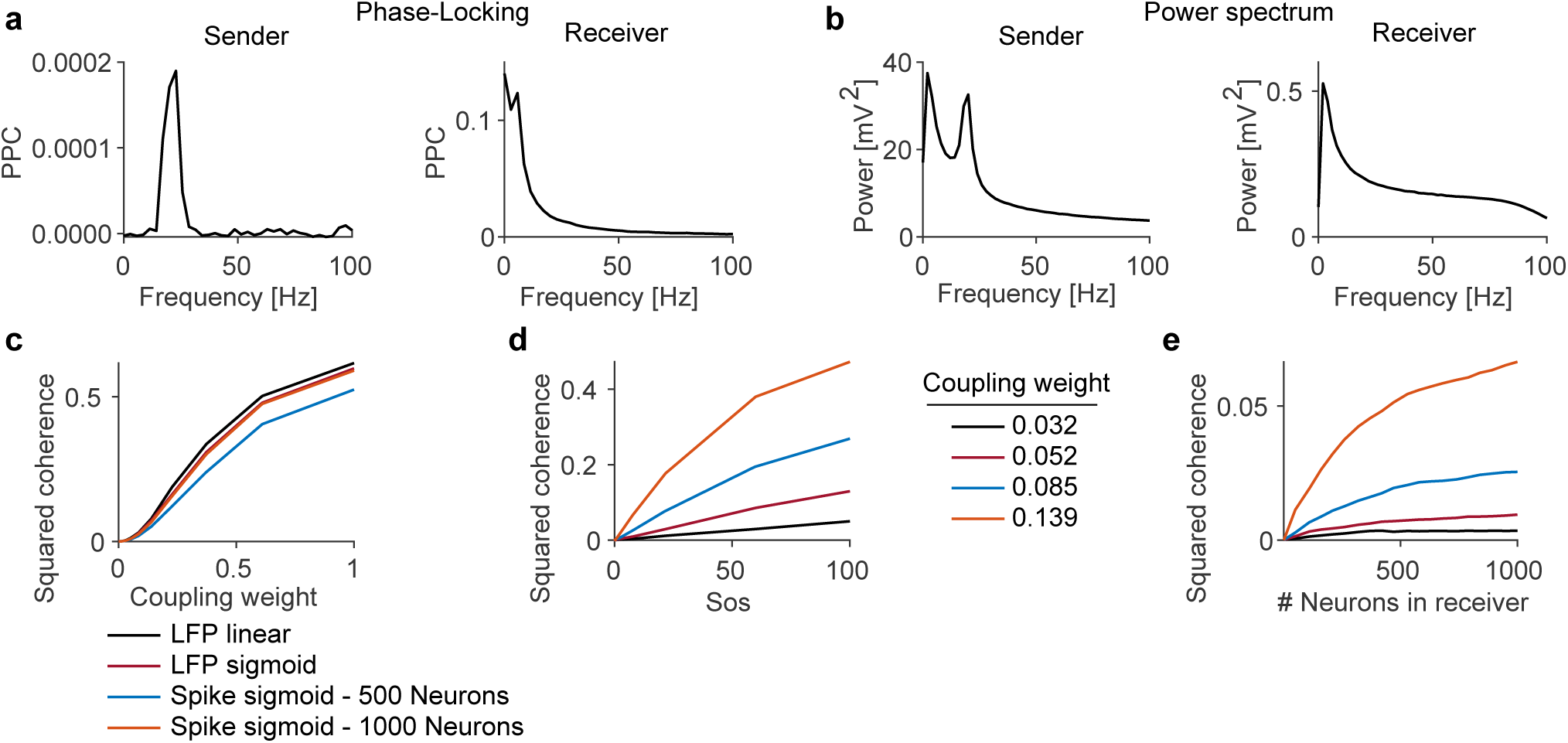
Neuron population activity distorted by sigmoidal input-output relationship approaches LFP-LFP coherence through synaptic mixing. Area 2 was modelled as a population of 1000 neurons modulated by the synthetic LFP signal of the extended synaptic mixing model (Figure 3c-f) according to inhomogeneous poisson process. **(a)** Phase-locking of receiver spikes to sender (left) and receiver (right) LFP. The phase-locking to the sender LFP signal shows a narrow-band peak at the modulation frequency. **(b)** Power spectra of both areas. The sender area follows the power spectrum expected from the model in Figure 3c-f. The power spectrum of the summed activity of neurons in the receiver area does not show a peak at frequency of the transmitted oscillatory signal. **(c)** For a high number of neurons the coherence between Area-1-LFP and Area-2 spiking approaches the LFP-LFP coherence. The LFP-LFP coherence (black) is generated with the synaptic mixing model from Figure 3c-f. In addition, we have generated an LFP-LFP model where the receiver signal depends sigmoidally on the transmitter signal. **(d)** The interareal coherence through sigmoidally modulated input-output relation of spiking shows dependecies on coupling and SOS similar to synaptic mixing. **(e)** Interareal coherence increases with the number of entrained neurons in the receiver. Depending on the coupling weight and the SOS the squared coherence saturates at a certain number of neurons.

**Figure S2:**
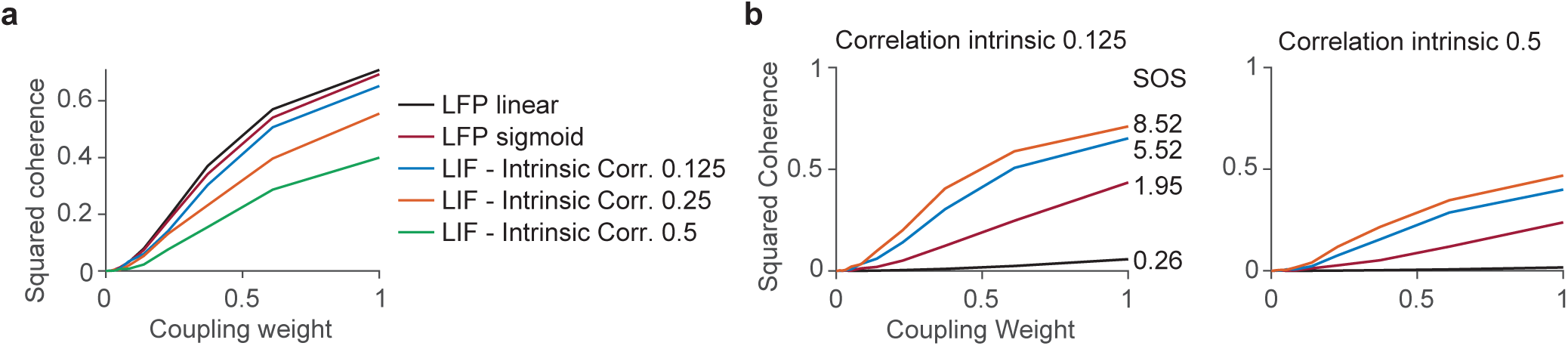
Dependence of coherence between Area-1 LFP and Area-2 spiking on connectivity, sender-oscillation-strength and number of neurons in Area 2. **(a)** Shown are three different kinds of models: (i) LFP-LFP coherence due to synaptic mixing. (ii) LFP-LFP coherence that results by passing the Area-2 signal through a sigmoid transformation, resulting in a decrease in coherence. (iii) Coherence between the summed population spiking activity in Area-2 and the Area-1 LFP. This shows a further decrease in coherence, especially when the intrinsic 1*/f* fluctuations in the receiver are highly correlated. The SOS for all models was 5.5. **(b)** Coherence between summed population spiking activity in the receiver and the sender LFP, for different values of SOS and coupling weight. Note similar dependences between LFP-spike and LFP-LFP coherence on SOS and coupling weight, but relatively higher LFP-LFP coherence due to synaptic mixing. Spike-LFP coherence is weaker when intrinsic correlations are stronger. Inter-areal coherence between Area-1 LFP and Area-2 spikes increases with the number of receiver neurons. The SOS is 5.5.

## Notes

### Competing Interest Statement

The authors have declared no competing interest.

### Summary of Updates

Added new Figure 6 on spike-LFP coherence, Added new Figure 3B, updated Discussion

## References

Abeles, M., 1982. Role of the cortical neuron: integrator or coincidence detector? Isr. Med. Assoc. J 18, 83.

Akam, T.E., Kullmann, D.M., 2012. Efficient “Communication through Coherence” Requires Oscillations Structured to Minimize Interference between Signals. PLOS Comp. Biol. 8, e1002760.

Babapoor-Farrokhran, S., Vinck, M., Womelsdorf, T., Everling, S., 2017. Theta and beta synchrony coordinate frontal eye fields and anterior cingulate cortex during sensorimotor mapping. Nature Communications, 13967.

Bastos, A.M., Vezoli, J., Bosman, C.A., Schoffelen, J.M., Oostenveld, R., Dowdall, J.R., De Weerd, P., Kennedy, H., Fries, P., 2015. Visual areas exert feedforward and feedback influences through distinct frequency channels. Neuron 85, 390–401.

Batista-Brito, R., Zagha, E., Ratliff, J.M., Vinck, M., 2018. Modulation of cortical circuits by top-down processing and arousal state in health and disease. Current opinion in neurobiology 52, 172–181.

Bernander, Ö., Koch, C., Douglas, R.J., 1994. Amplification and linearization of distal synaptic input to cortical pyramidal cells. J. Neurophysiol. 72, 2743–2753.

Bonnefond, M., Kastner, S., Jensen, O., 2017. Communication between brain areas based on nested oscillations. eneuro 4.

Börgers, C., Kopell, N.J., 2008. Gamma oscillations and stimulus selection. Neural Comput 20, 383–414.

Bosman, C., Schoffelen, J., Brunet, N., Oostenveld, R., Bastos, A., Womelsdorf, T., Rubehn, B., Stieglitz, T., De Weerd, P., Fries, P., 2012. Attentional stimulus selection through selective synchronization between monkey visual areas. Neuron 75, 875–888.

Bragin, A., Jandó, G., Nádasdy, Z., Hetke, J., Wise, K., Buzsáki, G., 1995. Gamma (40-100 hz) oscillation in the hippocampus of the behaving rat. J Neurosci 15, 47–60.

Bressler, S.L., 1995. Large-scale cortical networks and cognition. Brain Research Reviews 20, 288–304.

Bressler, S.L., Coppola, R., Nakamura, R., 1993. Episodic multiregional cortical coherence at multiple frequencies during visual task performance. Nature 366, 153–156.

Bressler, S.L., Richter, C.G., Chen, Y., Ding, M., 2006. Top-down cortical influences in visual expectation, in: IJCNN’06., pp. 188–194.

Brovelli, A., Ding, M., Ledberg, A., Chen, Y., Nakamura, R., Bressler, S.L., 2004. Beta oscillations in a large-scale sensorimotor cortical network: directional influences revealed by granger causality. Proc. Natl. Acad. Sci. U.S.A. 101, 9849–9854.

Brown, E.N., Barbieri, R., Ventura, V., Kass, R.E., Frank, L.M., 2002. The time-rescaling theorem and its application to neural spike train data analysis. Neural Computation 14, 325–346.

Brunet, N.M., Bosman, C.A., Vinck, M., Roberts, M., Oostenveld, R., Desimone, R., De Weerd, P., Fries, P., 2014. Stimulus repetition modulates gamma-band synchronization in primate visual cortex. Proc. Natl. Acad. Sci. U.S.A. 111, 3626–3631.

Buffalo, E.A., Fries, P., Landman, R., Buschman, T.J., Desimone, R., 2011. Laminar differences in gamma and alpha coherence in the ventral stream. Proc. Natl. Acad. Sci. U.S.A. 108, 11262–11267.

Burchell, T.R., Faulkner, H.J., Whittington, M.A., 1998. Gamma frequency oscillations gate temporally coded afferent inputs in the rat hippocampal slice. Neurosci Lett 255, 151–4.

Burns, S.P., Xing, D., Shapley, R.M., 2011. Is gamma-band activity in the local field potential of V1 cortex a “clock” or filtered noise? J. Neurosci. 31, 9658–9664.

Buschman, T.J., Miller, E.K., 2007. Top-down versus bottom-up control of attention in the prefrontal and posterior parietal cortices. Science 315, 1860–2.

Buzsáki, G., 2006. Rhythms of the Brain. Oxford University Press, USA.

Buzsáki, G., Anastassiou, C.A., Koch, C., 2012. The origin of extracellular fields and currents–EEG, ECoG, LFP and spikes. Nat. Rev. Neurosci. 13, 407–420.

Buzsáki, G., Draguhn, A., 2004. Neuronal oscillations in cortical networks. Science 304, 1926–1929.

Buzsáki, G., Schomburg, E.W., 2015. What does gamma coherence tell us about inter-regional neural communication? Nature neuroscience 18, 484–489.

Carmichael, J.E., Yuen, M.M., Van Der Meer, M.A., 2019. Piriform cortex provides a dominant gamma lfp oscillation in the anterior limbic system. BioRxiv, 861021.

Chalk, M., Herrero, J.L., Gieselmann, M.A., Delicato, L.S., Gotthardt, S., Thiele, A., 2010. Attention reduces stimulus-driven gamma frequency oscillations and spike field coherence in V1. Neuron 66, 114–25.

Chiu, C.Q., Lur, G., Morse, T.M., Carnevale, N.T., Ellis-Davies, G.C., Higley, M.J., 2013. Compartmentalization of gabaergic inhibition by dendritic spines. Science 340, 759–762.

Christensen, J.R., Larsen, K.B., Lisanby, S.H., Scalia, J., Arango, V., Dwork, A.J., Pakkenberg, B., 2007. Neocortical and hippocampal neuron and glial cell numbers in the rhesus monkey. The Anatomical Record: Advances in Integrative Anatomy and Evolutionary Biology: Advances in Integrative Anatomy and Evolutionary Biology 290, 330–340.

Colgin, L., Denninger, T., Fyhn, M., Hafting, T., Bonnevie, T., Jensen, O., Moser, M., Moser, E., 2009. Frequency of gamma oscillations routes flow of information in the hippocampus. Nature 462, 353–357.

Dann, B., Michaels, J.A., Schaffelhofer, S., Scherberger, H., 2016. Uniting functional network topology and oscillations in the fronto-parietal single unit network of behaving primates. Elife 5, e15719.

Donoghue, J.P., Sanes, J.N., Hatsopoulos, N.G., Gaal, G., 1998. Neural discharge and local field potential oscillations in primate motor cortex during voluntary movements. J Neurophysiol 79, 159–173.

Einevoll, G.T., Kayser, C., Logothetis, N.K., Panzeri, S., 2013. Modelling and analysis of local field potentials for studying the function of cortical circuits. Nat. Rev. Neurosci. 14, 770–785.

Engel, A.K., Fries, P., Singer, W., 2001. Dynamic predictions: oscillations and synchrony in top-down processing. Nat. Rev. Neurosci. 2, 704–716.

Ferro, D., van Kempen, J., Boyd, M., Panzeri, S., Thiele, A., 2020. Directed information exchange between cortical layers in macaque v1 and v4 and its modulation by selective attention. bioRxiv.

Fries, P., 2005. A mechanism for cognitive dynamics: neuronal communication through neuronal coherence. Trends Cogn. Sci. 9, 474–480.

Fries, P., 2009. Neuronal gamma-band synchronization as a fundamental process in cortical computation. Annu. Rev. Neurosci. 32, 209–224.

Fries, P., 2015. Rhythm for Cognition: Communication Through Coherence. Neuron 88, 220–235. 15334406.

Geweke, J., 1982. Measurement of linear dependence and feedback between multiple time series. Journal of the American Statistical Association 77, 304–313.

Gillespie, D.T., 1977. Exact stochastic simulation of coupled chemical reactions. The journal of physical chemistry 81, 2340–2361.

Gray, C., McCormick, D., 1996. Chattering cells: superficial pyramidal neurons contributing to the generation of synchronous oscillations in the visual cortex. Science 274, 109.

Gray, C.M., König, P., Engel, A.K., Singer, W., 1989. Oscillatory responses in cat visual cortex exhibit inter-columnar synchronization which reflects global stimulus properties. Nature 338, 334–337.

Gregoriou, G.G., Gotts, S.J., Zhou, H., Desimone, R., 2009. High-frequency, long-range coupling between prefrontal and visual cortex during attention. Science 324, 1207–1210.

Grothe, I., Neitzel, S.D., Mandon, S., Kreiter, A.K., 2012a. Switching neuronal inputs by differential modulations of gamma-band phase-coherence. J. Neurosci. 32, 16172–16180.

Grothe, I., Neitzel, S.D., Mandon, S., Kreiter, A.K., 2012b. Switching Neuronal Inputs by Differential Modulations of Gamma-Band Phase-Coherence. J. Neurosci. 32, 16172–16180.

Hagan, M.A., Dean, H.L., Pesaran, B., 2012. Spike-field activity in parietal area lip during coordinated reach and saccade movements. Journal of neurophysiology 107, 1275–1290.

Hahn, G., Bujan, A.F., Frégnac, Y., Aertsen, A., Kumar, A., 2014. Communication through resonance in spiking neuronal networks. PLoS Comput Biol 10, e1003811.

Han, Y., Kebschull, J.M., Campbell, R.A., Cowan, D., Imhof, F., Zador, A.M., Mrsic-Flogel, T.D., 2018. The logic of single-cell projections from visual cortex. Nature 556, 51–56.

Haufe, S., Nikulin, V.V., Nolte, G., 2012. Alleviating the influence of weak data asymmetries on granger-causal analyses, in: Latent Variable Analysis and Signal Separation. Springer, pp. 25–33.

Henrie, J.A., Shapley, R., 2005. LFP Power Spectra in V1 Cortex: The Graded Effect of Stimulus Contrast. J. Neurophysiol. 94, 479–490.

Hermes, D., Miller, K., Wandell, B., Winawer, J., 2015. Stimulus Dependence of Gamma Oscillations in Human Visual Cortex. Cereb. Cortex. 25, 2951–2959.

Johnson, P.B., Ferraina, S., Bianchi, L., Caminiti, R., 1996. Cortical networks for visual reaching: physiological and anatomical organization of frontal and parietal lobe arm regions. Cerebral cortex 6, 102–119.

Katzner, S., Nauhaus, I., Benucci, A., Bonin, V., Ringach, D.L., Carandini, M., 2009. Local origin of field potentials in visual cortex. Neuron 61, 35–41.

Kempter, R., Gerstner, W., Van Hemmen, J.L., Wagner, H., 1998. Extracting Oscillations: Neuronal Coincidence Detection with Noisy Periodic Spike Input. Neural Computation 10, 1987–2017.

van Kerkoerle, T., Self, M.W., Dagnino, B., Gariel-Mathis, M.A., Poort, J., van der Togt, C., Roelfsema, P.R., 2014. Alpha and gamma oscillations characterize feedback and feedforward processing in monkey visual cortex. Proc. Natl. Acad. Sci. U.S.A., 201402773.

Knoblich, U., Siegle, J.H., Pritchett, D.L., Moore, C.I., 2010. What do We Gain from Gamma? Local Dynamic Gain Modulation Drives Enhanced Efficacy and Efficiency of Signal Transmission. Front. Hum. Neurosci 04.

König, P., Engel, A.K., Roelfsema, P.R., Singer, W., 1995. How precise is neuronal synchronization? Neural Comput 7, 469–485.

Kreiter, A.K., 2006. How do we model attention-dependent signal routing? Neural networks 19, 1443–1444.

Lindén, H., Tetzlaff, T., Potjans, T.C., Pettersen, K.H., Grün, S., Diesmann, M., Einevoll, G.T., 2011. Modeling the Spatial Reach of the LFP. Neuron 72, 859–872.

Livingstone, M.S., 1996. Oscillatory firing and interneuronal correlations in squirrel monkey striate cortex. J. Neurophysiol. 75, 2467–2485.

Lowet, E., Roberts, M.J., Peter, A., Gips, B., De Weerd, P., 2017. A quantitative theory of gamma synchronization in macaque V1. eLife 6.

Luck, S.J., Chelazzi, L., Hillyard, S.A., Desimone, R., 1997. Neural mechanisms of spatial selective attention in areas v1, v2, and v4 of macaque visual cortex. Journal of neurophysiology 77, 24–42.

Lund, J.S., Angelucci, A., Bressloff, P.C., 2003. Anatomical substrates for functional columns in macaque monkey primary visual cortex. Cereb. Cortex. 13, 15–24.

Luppino, G., Murata, A., Govoni, P., Matelli, M., 1999. Largely segregated parietofrontal connections linking rostral intraparietal cortex (areas aip and vip) and the ventral premotor cortex (areas f5 and f4). Experimental Brain Research 128, 181–187.

Lur, G., Vinck, M.A., Tang, L., Cardin, J.A., Higley, M.J., 2016. Projection-Specific Visual Feature Encoding by Layer 5 Cortical Subnetworks. Cell Reports 14, 2538–2545.

Markov, N., Misery, P., Falchier, A., Lamy, C., Vezoli, J., Quilodran, R., Gariel, M., Giroud, P., Ercsey-Ravasz, M., Pilaz, L., et al., 2011. Weight consistency specifies regularities of macaque cortical networks. Cerebral cortex 21, 1254–1272.

Markov, N.T., Vezoli, J., Chameau, P., Falchier, A., Quilodran, R., Huissoud, C., Lamy, C., Misery, P., Giroud, P., Ullman, S., et al., 2014. Anatomy of hierarchy: Feedforward and feedback pathways in macaque visual cortex. Journal of Comparative Neurology 522, 225–259.

McGinley, M.J., Vinck, M., Reimer, J., Batista-Brito, R., Zagha, E., Cadwell, C.R., Tolias, A.S., Cardin, J.A., McCormick, D.A., 2015. Waking State: Rapid Variations Modulate Neural and Behavioral Responses. Neuron 87, 1143–1161.

Mejias, J.F., Murray, J.D., Kennedy, H., Wang, X.J., 2016. Feedforward and feedback frequency-dependent interactions in a large-scale laminar network of the primate cortex. Science advances 2, e1601335.

Michalareas, G., Vezoli, J., van Pelt, S., Schoffelen, J.M., Kennedy, H., Fries, P., 2016. Alpha-Beta and Gamma Rhythms Subserve Feedback and Feedforward Influences among Human Visual Cortical Areas. Neuron 89, 384–397. 15334406.

Miller, E.K., Wilson, M.A., 2008. All my circuits: using multiple electrodes to understand functioning neural networks. Neuron 60, 483–8.

Mitzdorf, U., 1985. Current source-density method and application in cat cerebral cortex: investigation of evoked potentials and EEG phenomena. Physiol. Rev. 65, 37–100.

Montgomery, S.M., Buzsáki, G., 2007. Gamma oscillations dynamically couple hippocampal ca3 and ca1 regions during memory task performance. Proc. Natl. Acad. Sci. U.S.A. 104, 14495–14500.

Murray, J.D., Bernacchia, A., Freedman, D.J., Romo, R., Wallis, J.D., Cai, X., Padoa-Schioppa, C., Pasternak, T., Seo, H., Lee, D., et al., 2014. A hierarchy of intrinsic timescales across primate cortex. Nature neuroscience 17, 1661.

Murthy, V.N., Fetz, E.E., 1996. Oscillatory activity in sensorimotor cortex of awake monkeys: synchronization of local field potentials and relation to behavior. J. Neurophysiol. 76, 3949–3967.

Nawrot, M.P., Boucsein, C., Rodriguez Molina, V., Riehle, A., Aertsen, A., Rotter, S., 2008. Measurement of variability dynamics in cortical spike trains. Journal of Neuroscience Methods 169, 374–390.

Nicolelis, M.A., Baccala, L.A., Lin, R., Chapin, J.K., 1995. Sensorimotor encoding by synchronous neural ensemble activity at multiple levels of the somatosensory system. Science 268, 1353–1358.

Nolte, G., Bai, O., Wheaton, L., Mari, Z., Vorbach, S., Hallett, M., 2004. Identifying true brain interaction from EEG data using the imaginary part of coherency. Clin Neurophysiol 115, 2292–2307.

Nunez, P.L., Srinivasan, R., 2006. Electric fields of the brain: the neurophysics of EEG? Oxford University Press.

Olcese, U., Bos, J.J., Vinck, M., Lankelma, J.V., van Mourik-Donga, L.B., Schlumm, F., Pennartz, C.M., 2016. Spike-based functional connectivity in cerebral cortex and hippocampus: loss of global connectivity is coupled to preservation of local connectivity during non-rem sleep. Journal of Neuroscience 36, 7676–7692.

Onorato, I., Neuenschwander, S., Hoy, J., Lima, B., Rocha, K.S., Broggini, A.C., Uran, C., Spyropoulos, G., Klon-Lipok, J., Womelsdorf, T., Fries, P., Niell, C., Singer, W., Vinck, M., 2020. A distinct class of bursting neurons with strong gamma synchronization and stimulus selectivity in monkey v1. Neuron 105, 180–197.

Oostenveld, R., Fries, P., Maris, E., Schoffelen, J.M., 2011. FieldTrip: Open source software for advanced analysis of MEG, EEG, and invasive electrophysiological data. Comput Intell Neurosci 2011, 156869.

Palmigiano, A., Geisel, T., Wolf, F., Battaglia, D., 2017. Flexible information routing by transient synchrony. Nature neuroscience 20, 1014.

Parabucki, A., Lampl, I., 2017. Volume conduction coupling of whisker-evoked cortical lfp in the mouse olfactory bulb. Cell reports 21, 919–925.

Pesaran, B., Vinck, M., Einevoll, G., Sirota, A., Fries, P., Siegel, M., Truccolo, W., Schroeder, C., Srinivasan, R., 2018. Investigating large-scale brain dynamics using field potential recordings: analysis and interpretation. Nat. Neurosci..

Peter, A., Uran, C., Klon-Lipok, J., Roese, R., Van Stijn, S., Barnes, W., Dowdall, J.R., Singer, W., Fries, P., Vinck, M., 2019. Surface color and predictability determine contextual modulation of v1 firing and gamma oscillations. eLife 8, e42101.

Phillips, J.M., Vinck, M., Everling, S., Womelsdorf, T., 2014. A long-range fronto-parietal 5-to 10-hz network predicts “top-down” controlled guidance in a task-switch paradigm. Cerebral Cortex 24, 1996–2008.

Pike, F., Goddard, R., Suckling, J., Ganter, P., Kasthuri, N., Paulsen, O., 2000. Distinct frequency preferences of different types of rat hippocampal neurones in response to oscillatory input currents. The Journal of Physiology 529, 205.

Powanwe, A.S., Longtin, A., 2019. Determinants of brain rhythm burst statistics. Scientific Reports 9, 1–23.

Ray, S., Maunsell, J.H., 2015. Do gamma oscillations play a role in cerebral cortex? Trends in cognitive sciences 19, 78–85.

Richter, C.G., Coppola, R., Bressler, S.L., 2018. Top-down beta oscillatory signaling conveys behavioral context in early visual cortex. Sci. Rep. 8, 6991.

Roberts, M.J., Lowet, E., Brunet, N.M., Ter Wal, M., Tiesinga, P., Fries, P., De Weerd, P., 2013. Robust Gamma Coherence between Macaque V1 and V2 by Dynamic Frequency Matching. Neuron 78, 523–536.

Salazar, R., Dotson, N., Bressler, S., Gray, C., 2012a. Content-specific frontoparietal synchronization during visual working memory. Science 338, 1097–1100.

Salazar, R.F., Dotson, N.M., Bressler, S.L., Gray, C.M., 2012b. Content-Specific Fronto-Parietal Synchronization During Visual Working Memory. Science 338, 1097–1100.

Saleem, A.B., Lien, A.D., Krumin, M., Haider, B., Rosón, M.R., Ayaz, A., Reinhold, K., Busse, L., Carandini, M., Harris, K.D., 2017. Subcortical Source and Modulation of the Narrowband Gamma Oscillation in Mouse Visual Cortex. Neuron 93, 315–322.

Salinas, E., Sejnowski, T.J., 2001. Correlated neuronal activity and the flow of neural information. Nat Rev Neurosci 2, 539–50.

Scherberger, H., Jarvis, M.R., Andersen, R.A., 2005. Cortical local field potential encodes movement intentions in the posterior parietal cortex. Neuron 46, 347–354.

Schomburg, E.W., Fernández-Ruiz, A., Mizuseki, K., Berényi, A., Anastassiou, C.A., Koch, C., Buzsáki, G., 2014. Theta phase segregation of input-specific gamma patterns in entorhinal-hippocampal networks. Neuron 84, 470–485.

Sejnowski, T.J., Paulsen, O., 2006. Network oscillations: emerging computational principles. J. Neurosci. 26, 1673–1676.

Siegle, J.H., Jia, X., Durand, S., Gale, S., Bennett, C., Graddis, N., Heller, G., Ramirez, T.K., Choi, H., Luviano, J.A., et al., 2019. A survey of spiking activity reveals a functional hierarchy of mouse corticothalamic visual areas. bioRxiv, 805010.

Singer, W., Gray, C.M., 1995. Visual feature integration and the temporal correlation hypothesis. Annu. Rev. Neurosci. 18, 555–586.

Sirota, A., Montgomery, S., Fujisawa, S., Isomura, Y., Zugaro, M., Buzsáki, G., 2008a. Entrainment of neocortical neurons and gamma oscillations by the hippocampal theta rhythm. Neuron 60, 683–697.

Sirota, A., Montgomery, S., Fujisawa, S., Isomura, Y., Zugaro, M., Buzsáki, G., 2008b. Entrainment of Neocortical Neurons and Gamma Oscillations by the Hippocampal Theta Rhythm. Neuron 60, 683–697.

Spyropoulos, G., Dowdall, J.R., Schölvinck, M.L., Bosman, C.A., Lima, B., Peter, A., Onorato, I., Klon-Lipok, J., Roese, R., Neuenschwander, S., Wolf, S., Vinck, M., Fries, P., 2020. Spontaneous variability in gamma dynamics described by a linear harmonic oscillator driven by noise. bioRxiv, 793729.

Tiesinga, P., Sejnowski, T., 2010. Mechanisms for phase shifting in cortical networks and their role in communication through coherence. Front. Hum. Neurosci 4.

Trongnetrpunya, A., Nandi, B., Kang, D., Kocsis, B., Schroeder, C.E., Ding, M., 2016. Assessing granger causality in electrophysiological data: removing the adverse effects of common signals via bipolar derivations. Frontiers in systems neuroscience 9, 189.

Varela, F., Lachaux, J.P., Rodriguez, E., Martinerie, J., 2001. The brainweb: phase synchronization and large-scale integration. Nat. Rev. Neurosci. 2, 229–239.

Vinck, M., Battaglia, F.P., Womelsdorf, T., Pennartz, C., 2012. Improved measures of phase-coupling between spikes and the Local Field Potential. J Comput Neurosci 33, 53–75.

Vinck, M., Bos, J.J., Mourik-Donga, V., Laura, A., Oplaat, K.T., Klein, G.A., Jackson, J.C., Gentet, L.J., Pennartz, C., 2016. Cell-type and state-dependent synchronization among rodent somatosensory, visual, perirhinal cortex, and hippocampus ca1. Frontiers in systems neuroscience 9, 187.

Vinck, M., Bosman, C.A., 2016. More gamma more predictions: Gamma-synchronization as a key mechanism for efficient integration of classical receptive field inputs with surround predictions. Front Syst Neurosci 10, 35.

Vinck, M., Huurdeman, L., Bosman, C.A., Fries, P., Battaglia, F.P., Pennartz, C.M., Tiesinga, P.H., 2015. How to detect the granger-causal flow direction in the presence of additive noise? Neuroimage 108, 301–318.

Vinck, M., Oostenveld, R., van Wingerden, M., Battaglia, F., Pennartz, C.M., 2011. An improved index of phase-synchronization for electrophysiological data in the presence of volume-conduction, noise and sample-size bias. Neuroimage 55, 1548–1565.

Vinck, M., van Wingerden, M., Womelsdorf, T., Fries, P., Pennartz, C.M., 2010. The pairwise phase consistency: a bias-free measure of rhythmic neuronal synchronization. Neuroimage 51, 112–122.

Vinck, M., Womelsdorf, T., Buffalo, E.A., Desimone, R., Fries, P., 2013a. Attentional modulation of cell-class-specific gamma-band synchronization in awake monkey area V4. Neuron 80, 1077–1089.

Vinck, M., Womelsdorf, T., Fries, P., 2013b. Gamma-band synchronization and information transmission, in: Quiroga-Quian, R., Panzeri, S. (Eds.), Principles of Neural Coding. CRC Press.

Volgushev, M., Chistiakova, M., Singer, W., 1998. Modification of discharge patterns of neocortical neurons by induced oscillations of the membrane potential. Neuroscience 83, 15–25.

Von Stein, A., Sarnthein, J., 2000. Different frequencies for different scales of cortical integration: from local gamma to long range alpha/theta synchronization. International Journal of Psychophysiology 38, 301–313.

Wallace, E., Benayoun, M., Van Drongelen, W., Cowan, J.D., 2011. Emergent oscillations in networks of stochastic spiking neurons. Plos one 6.

Wilson, H.R., Cowan, J.D., 1972. Excitatory and inhibitory interactions in localized populations of model neurons. Biophys. J. 12, 1–24.

Xing, D., Yeh, C.I., Burns, S., Shapley, R.M., 2012. Laminar analysis of visually evoked activity in the primary visual cortex. Proc. Natl. Acad. Sci. U.S.A. 109, 13871–13876.

Zandvakili, A., Kohn, A., 2015. Coordinated Neuronal Activity Enhances Corticocortical Communication. Neuron 87, 827–839.

Zeitler, M., Fries, P., Gielen, S., 2006. Assessing neuronal coherence with single-unit, multi-unit, and local field potentials. Neural Comput 18, 2256–2281.

